# Developmental asymmetries in learning to adjust to cooperative and uncooperative environments

**DOI:** 10.1101/2020.07.29.226332

**Authors:** Bianca Westhoff, Lucas Molleman, Essi Viding, Wouter van den Bos, Anna C. K. van Duijvenvoorde

## Abstract

Learning to successfully navigate social environments is a critical developmental goal, predictive of long-term wellbeing. However, little is known about how people learn to adjust to different social environments, and how this behaviour emerges across development. Here, we use a series of economic games to assess how children, adolescents, and young adults learn to adjust to social environments that differ in their level of cooperation (i.e., trust and coordination). Our results show an asymmetric developmental pattern: adjustment requiring uncooperative behaviour remains constant across adolescence, but adjustment requiring cooperative behaviour improves markedly across adolescence. Behavioural and computational analyses reveal that age-related differences in this social learning are shaped by age-related differences in the degree of inequality aversion and in the updating of beliefs about others. Our findings point to early adolescence as a phase of rapid change in cooperative behaviours, and highlight this as a key developmental window for interventions promoting well-adjusted social behaviour.

## Introduction

Humans have evolved in a highly social environment in which they continuously make decisions about how to engage with others. Well-adjusted social behaviour requires individuals to learn whom they can trust and cooperate with. We typically trust others whom we expect to reciprocate that trust in the future and beliefs about others’ trustworthiness are updated through everyday experiences. For example, if a friend violates our trust, this calls for an adjustment of our belief in their trustworthiness. This may not happen on the first violation, but if this friend continues their untrustworthy behaviour, the friendship is unlikely to survive. Adjusting our beliefs based on outcomes of social interactions enables decision making that matches the situation and can be critical for successful navigation of the social world. In line with this notion, well-adjusted social behaviour has been linked to positive developmental trajectories (e.g., in health, education, and social development), and is important for long-term mental health ^1–4^. With the rise of complex social worlds (both online and offline), learning about and adjusting to different social environments may be more important than ever. Yet little is known about the factors that underlie learning and adjusting in social environments, and how these skills manifest across adolescence.

Mounting evidence suggests that adolescence – the period between childhood and young adulthood – is a life phase in which learning and flexible behaviour mature rapidly (see e.g., ^1,3,5,6^). Moreover, adolescence is marked by a social reorientation: individuals start to form larger peer groups, peers gain in importance compared with parents, and social interactions become more complex ^7–10^. Adolescence is, therefore, an important life phase for developing well-adjusted social behaviour, with cooperative and uncooperative behaviours becoming more salient as adolescents deal with their social environments more independently. Developmental research into social decision-making has shown that from an early age, children trust others and recognize that investing in others can lead to mutual benefits ^11^. Cooperative behaviours, such as trust and prosocial behaviour, are thought to continuously increase during adolescence ^5,12–14^ (but see ^15^). A more nuanced view is that adolescents do not show more cooperative behaviours per se, but instead increasingly tailor their behaviour to the social environment. For example, when undertaking prosocial actions they increasingly differentiate between friends and strangers^16–18^. Also, adolescents learn to adjust to interaction partners that differ in their level of trustworthiness ^19^, something that children find difficult ^11^.

Theoretically, decisions to cooperate can be based on at least three distinctive factors, each of which could differ between individuals and could contribute to developmental differences in adjusting behaviour in social contexts: (1) social preferences, (2) prior expectations, and (3) updating of expectations. First, *social preferences* refer to individuals caring about relative outcomes, i.e. disliking having either a better or worse outcome than others ^20^. Across childhood, there is a decrease in the preference of avoiding getting less than others (i.e., disadvantageous inequality aversion) and an increase in the preference of avoiding getting *more* than others (i.e., advantageous inequality aversion ^21^; and see ^22^ for a review). Although evidence suggests that social preferences continue to develop across adolescence (e.g., ^23^), little is known about how they impact learning in social environments. For instance, high levels of disadvantageous inequality aversion could prevent cooperative behaviour due to a fear of getting less than others, even if there is a relatively strong expectation that others will cooperate.

Second, *prior expectations* (i.e., descriptive norms, the perceptions of what most people do; ^24–26^) inform decision-making by generating predictions about the behaviour of others (e.g., I will cooperate if this person is likely to reciprocate). Individual differences in initial expectations about others may lead to individual differences in choices ^27^, and it is conceivable that different age groups have varying prior expectations. However, prior expectations have hardly been studied in developmental populations, despite being important determinants of cooperative behaviours.

Third, adjusting behaviour requires expectations to be *updated* in response to new information. Updating of expectations in social environments can be captured by reinforcement learning (RL) models (e.g., ^28–30^), in which learning is driven by differences between expected and received rewards (i.e., prediction errors). Adolescence is characterized by substantial improvements in flexible learning and quick adaptation to novel non-social contexts (^31–33^); whether this extends to the social domain, however, is still unclear (but see ^34^).

Here we examine experimentally how children, adolescents, and adults adjust to social environments that differ in their level of cooperation, and aim to provide a mechanistic explanation by evaluating the role of social preferences, prior expectations, and expectation updating. To achieve this goal, we deployed a set of economic games, together with behavioural analyses and computational reinforcement learning modelling. Our cross-sectional sample spanned from late childhood into early adulthood (8 to 23 years old, *N*=244). Participants played age-appropriate versions of two well-studied incentivized economic games: A Trust Game (Figure 1b) and a Coordination Game (Figure 1d). These two games involve key types of cooperative behaviours: trust and coordination. Trust is key for mutually beneficial cooperation to be initiated and sustained (e.g., ^12,19,35^), and for achieving beneficial outcomes for all interaction partners involved. Yet, trust also creates a hazard of being betrayed. Similarly, coordinating one’s behaviour with others is often critical for collective welfare, even though outcomes may not always equally benefit all interaction partners ^36,37^.

**Figure 1.**
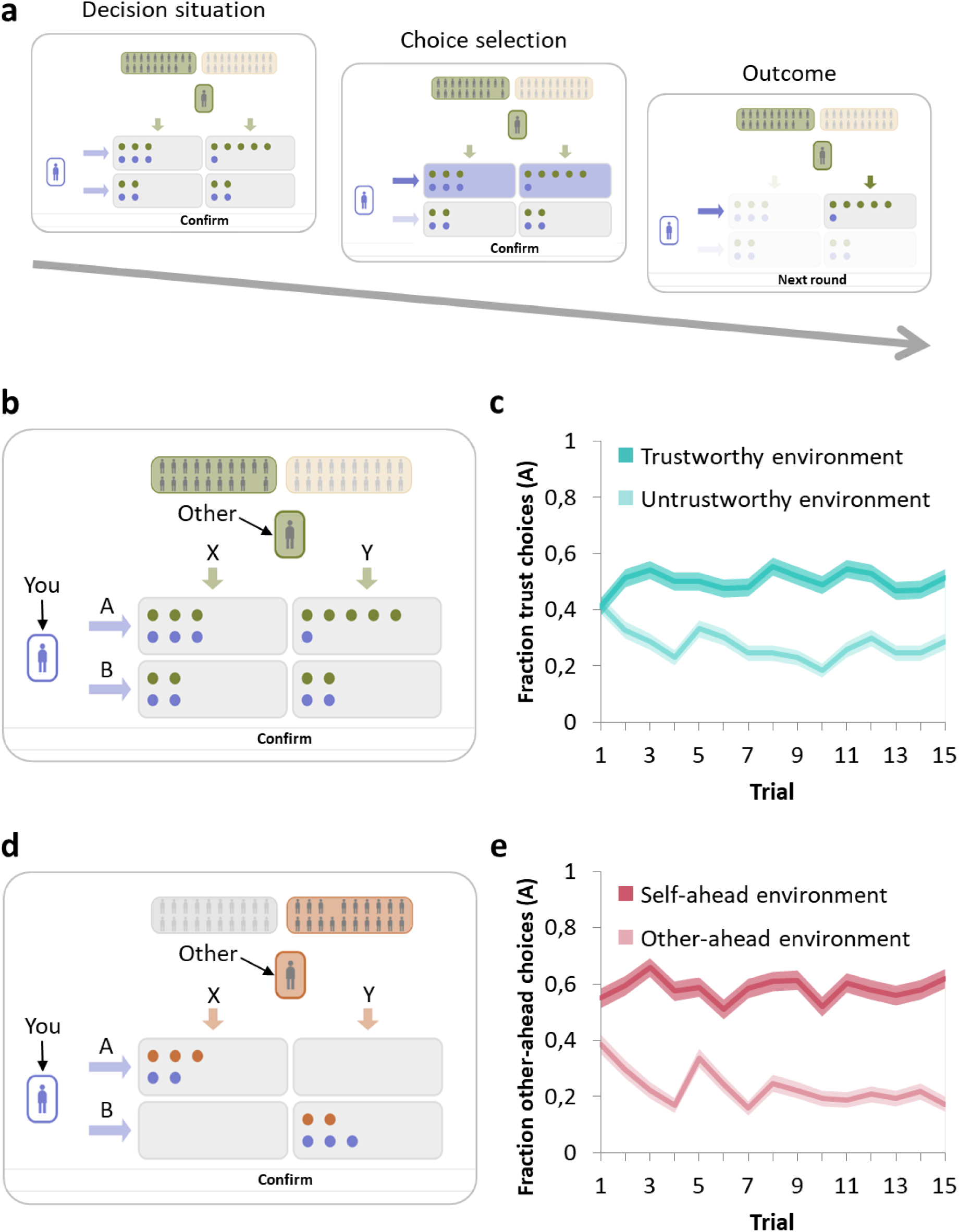
Task assessing learning to adjust to cooperative and uncooperative social environments. **a**, Example trial. The participant (purple stick figure on the left) can choose between the top and bottom row of boxes (A or B). After choice selection, the participant is shown the pre-recorded choice (X or Y) of the other player (grey stick figure on the top). The background colour of the other player indicates to which of the two environments they belong. The combined choices of the participant and the other player determine the monetary outcome for both players (number of dots in their corresponding colour). **b**, In the Trust Game, participants interact with players from a ‘trustworthy’ environment (who tend to choose X) or an ‘untrustworthy’ environment (who tend to choose Y). Participants’ own monetary payoffs are maximized by choosing to trust (choose A) a player from a trustworthy environment, and to withhold trust (choose B) from a player from the untrustworthy environment. In this trust-game setup only disadvantageous inequality aversion may play a role in decision making **c**, Participant choices over trials per social environment, pooled across all participants. Over the course of the game, participants adjusted their choices by directing their trust towards players from the trustworthy environment, and away from players from the untrustworthy environment (N=244). **d**, In the Coordination Game, participants’ monetary payoffs are also maximised by matching the choices of their co-players. Again, social environments differ in their prevalence of (non)cooperation, reflected by players’ tendencies to choose either X or Y. Coordinating on either of the outcomes (A,X) or (B,Y) will lead to positive outcomes. Respectively, participants’ own monetary payoffs are maximized by choosing to put themselves behind (choose A) when confronted with a player from the self-ahead environment, and to put themselves ahead (choose B) when confronted with a player from the other-ahead environment. In this game both advantageous and disadvantageous inequality aversion may play a role in social decision-making. **e**, Participant choices over trials per social environment, pooled across all participants. Overall, participants learned to coordinate with players from both environments. (N=202). Shaded areas in panels c and e represent standard errors of the mean (s.e.m.).

The two games consisted of repeated one-shot interactions, in which both players had to choose between two options. In each trial, they encountered one new anonymous player from either a Cooperative environment or an Uncooperative environment. The decisions of these players had been recorded in a previous session with age-matched unfamiliar others (see *Methods, pre-test*). Participants were explained that between environments, players could differ in their tendency to choose X (see Figure 1b and 1d). To maximise their earnings, participants had to learn which environment was cooperative and which was uncooperative over the course of the game, and adjust their choices accordingly. The social environments in these games were probabilistic, as cooperative behaviours were displayed by 73% of the players in the cooperative environments, and by 27% of the players in the uncooperative environments.

Participants also played an iterative Ultimatum Game (UG) and Dictator Game (DG), which allowed us to estimate participants’ social preferences (i.e., advantageous and disadvantageous inequality aversion; see *Methods*). We separately assessed participants’ prior expectations of the behaviour of others before the start of the Trust Game and Coordination Game (see *Methods*). Furthermore, we used computational reinforcement-learning models ^38^ to model the updating of expectations between interactions. In these models, the learning rate quantifies how much an expectation violation modifies our subsequent expectations and consequently our decision-making. We allowed learning rates to decay over the course of the games because we expected that most of the learning about the environments would happen in the first set of trials. After that behaviour would stabilize, provided the environments did not change their behaviour (for more on learning rates and environmental stability see ^39–41^). We extended these reinforcement learning models to account for the measured prior expectations and social preferences ^28^, and compared the parameters of these models across age cohorts (see *Methods*).

We hypothesized that participants would be able to learn to adjust their behaviours to social environments differing in their level of (non)cooperation, but that across adolescence this ability would improve rapidly. We expected that these developmental differences could be explained by a combination of (1) social preferences (i.e., age-related changes in levels of advantageous and disadvantageous inequality aversion), (2) prior expectations (i.e., age-related changes in expectations about others’ trustworthiness and tendencies to prioritise their own payoffs over those of others) and (3) updating of expectations (i.e., age-related changes in learning rates).

## Results

### Learning to adjust to cooperative and uncooperative social environments across different ages

First, we examined decisions over the course of the games to assess whether children, adolescents, and young adults adjust their behaviour to different social environments with different levels of cooperation. For this, we used the Trust Game in which participants maximized their monetary outcomes by trusting trustworthy others and withhold trust from untrustworthy others (Figure 1b), and the Coordination Game in which participants maximized their outcomes by coordinating with the response of the others placing themselves ahead or behind. We performed a binomial generalized linear mixed model (GLMM) per game on participants’ binary choices, including social preferences and prior estimations of others behaviour (see *Methods*).

For the Trust Game, results indicated an accelerated change in adolescence in which people differentiated more between the trustworthy and untrustworthy environment (environment x age linear, *B* = -0.3, *P* < 0.001; environment x age quadratic, *B* = 0.207, *P* = 0.014; N=244; see Table S1 for full statistical analysis; Figure 2a). Post-hoc tests per social environment showed that trusting the trustworthy others increased rapidly between ages 8-11 and 15-17 (age linear, *B* = -0.374, *P* = 0.009; age quadratic, *B* = 0.322, *P* = 0.018). In contrast, adjusting to untrustworthy others improved slightly, and monotonically across adolescence (age linear, *B* = 0.229, *P* = 0.037).

**Figure 2.**
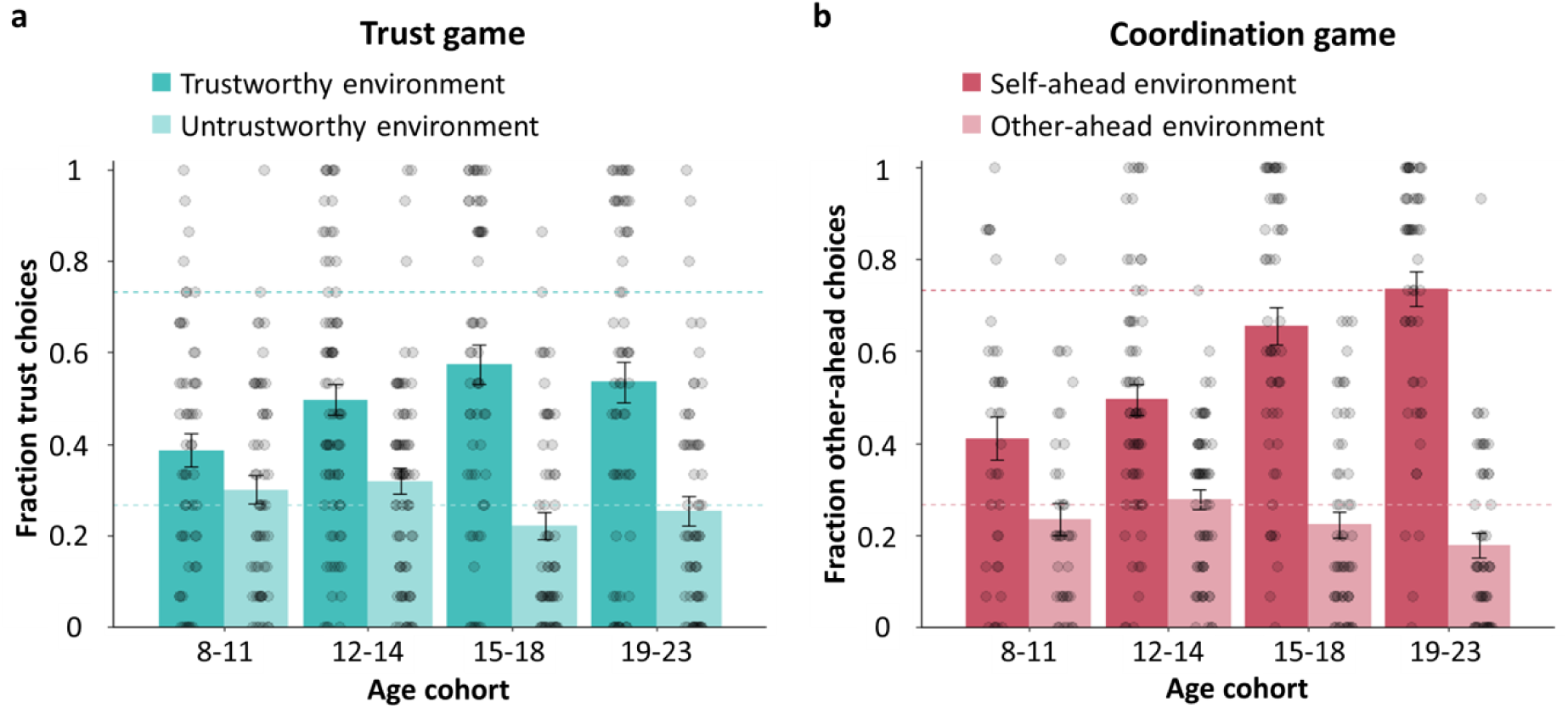
Developmental asymmetries in adjusting to cooperative and uncooperative social environments. Decisions per age cohort are shown per social environment in the (**a**) Trust Game (N = 244) and (**b**) Coordination Game (N = 202). Across age cohorts, there is a pronounced increase in Cooperative choices (adjusting to a trustworthy and a self-ahead environment), whereas Uncooperative choices (adjusting to an untrustworthy and self-behind environment) were relatively stable across age. Dashed lines in a and b indicate the reinforcement rates (i.e., fraction of 0.73 and 0.23) for each social environment. Note that we grouped participants for illustration purposes only and age was treated as a continuous variable in all analyses. Error bars represent s.e.m.

For the Coordination Game, results again indicated that with age, people differentiated more between the self-ahead and other-ahead environment (environment x age linear, *B* = -0.446, *P* < 0.001; N = 202; see Table S3 for full statistical analysis; Figure 2b). Post-hoc tests per social environment showed that optimally coordinating to the other-ahead environment (participant having fewer points than the other player) increased across adolescence (age linear, *B* = -0.446, *P* < 0.001). However, coordinating to the self-ahead environment (participant having more points than the other player) did not change with age; participants from all age cohorts adjusted quickly to this environment.

### Social preferences and prior expectations

Social preferences (advantageous and disadvantageous inequality aversion) and prior expectations of others’ behaviour are features that may account for age-related changes in learning to adjust to different social environments. Before further testing their relation to behaviour in the Trust Game and Coordination Game, we first examined the age-related changes in these parameters. Robust linear regression analyses (5000 bootstraps) indicated that only disadvantageous inequality aversion changed across age (Figure 3a-3d). Specifically, older participants were less averse to being behind than younger participants (age linear, B = -.127, β = -0.324, *P* < 0.001, 95% CI [-0.172, -0.083], N = 244). We did not observe significant age-related change for advantageous inequality aversion (age linear, B = .01, β = 0.123, *P* = 0.071, 95% CI [-0.001, 0.021], N = 202), nor for prior expectations of others’ trustworthiness (age linear, *P* = 0.462; N = 245) or for prior expectations of others’ tendency to have more than the other (age linear, *P* = 0.478, N = 245).

**Figure 3.**
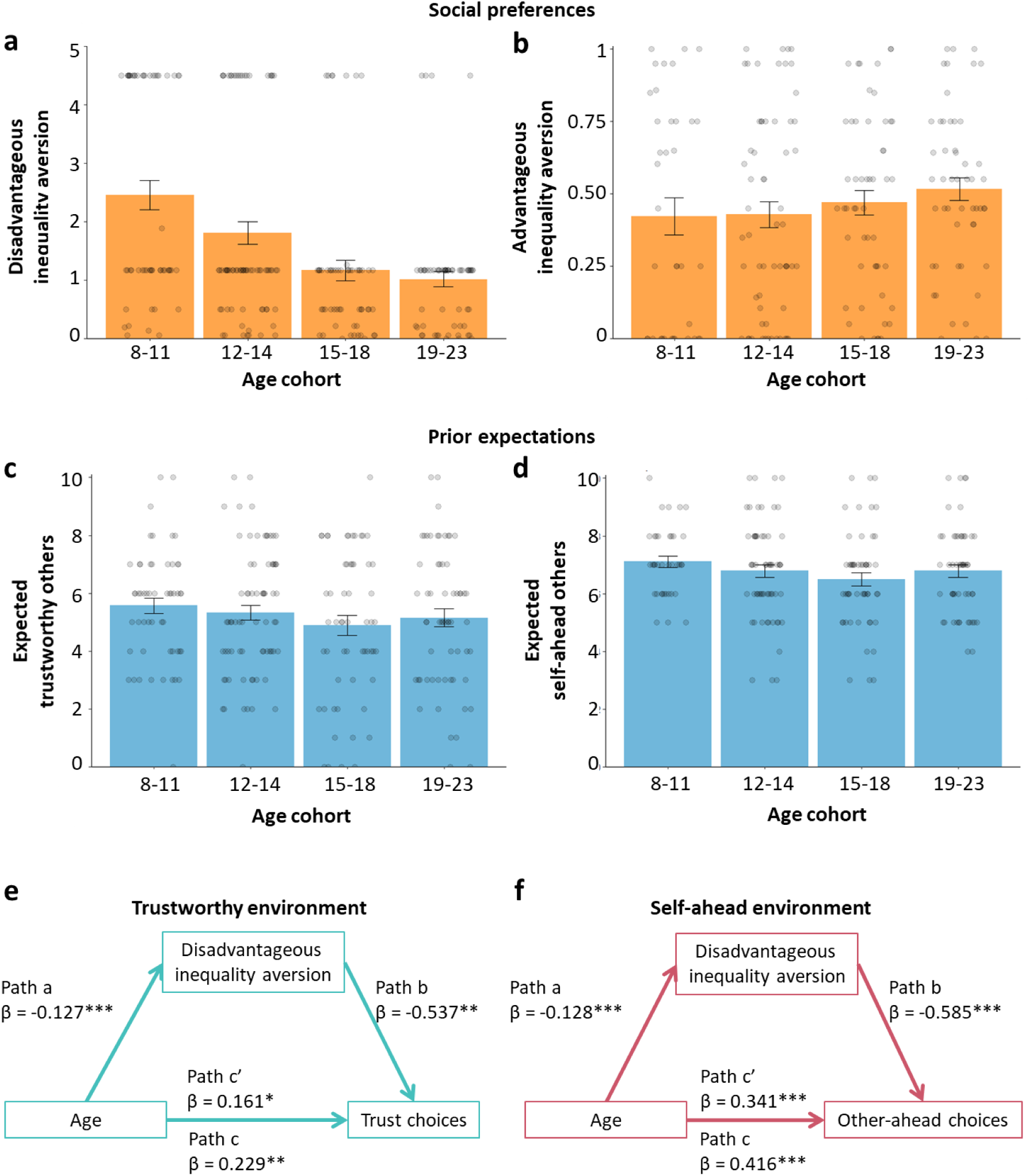
Social preferences and prior expectations across age cohorts. Social preferences and prior expectations are features that may account for choices in the games. **a**, Estimated disadvantageous inequality aversion and **b**, estimated advantageous inequality aversion were used as social preference measures. **c**, Prior expectations for the Trust Game, with higher values indicating greater expectations that others are trustworthy. **d**, Prior expectations for the Coordination Game, with higher values indicating greater expectations that others will choose to put themselves behind. In panels a-d, error bars show s.e.m. **e**, Mediation model for the effect of age on trust behaviour towards players from the trustworthy environment, via disadvantageous inequality aversion. **f**, Mediation model for the effect of age on coordination behaviour towards players from the other-ahead environment, via disadvantageous inequality aversion. Note in mediation models: c = total effect, c’ = direct effect; values are standardized regression coefficients of direct effects, and asterisks indicate significance levels (\**P* < 0.05, \*\**P* < 0.01, \*\*\**P* < 0.001).

In a binomial GLMM analyses, advantageous and disadvantageous inequality aversion was related to choices in the games (Tables S1 and S3). Greater disadvantageous inequality aversion was associated with overall less trusting choices (*B* = 0.194, *P* < 0.027) and with less other-ahead choices (*B* = 0.184, *P* < 0.007). In addition, greater advantageous inequality aversion was associated with more other-ahead choices (*B* = -0.162, *P* < 0.014). In contrast, prior expectations were not related to choices in both games.

To better understand what drives the age-related change in learning to adjust to the social environments differing in their level of cooperation, we ran a mediation analysis per game. Specifically, we examined whether the age-related changes in disadvantageous inequality aversion explained the observed increase in cooperative behaviour across development controlling for individual’s level of uncooperative behaviour (Figure 3e; Figure 3f). We found that the improvement across age in adjusting to the trustworthy environment (β = 0.161, *P* =.040) was partly explained by the age-related decrease in disadvantageous inequality aversion (indirect effect = 0.068, *SE* = 0.027 CI [0.017, 0.125]). That is, older participants showed lower levels of disadvantageous inequality aversion (β = -0.127, *P* < 0.001), which in turn resulted in more trust choices (β = -0.537, *P* = 0.007; Figure 3e). This mediation analysis for the Coordination Game showed a similar effect of disadvantageous inequality aversion partly explaining the age-related change in cooperative behaviour. That is, older participants showed lower levels lower levels of disadvantageous inequality aversion (β = -0.128, *P* < 0.001), which in turn resulted in less cooperative (other-ahead) choices (β = -0.585, *P* = < 0.001; Figure 3f). Note that this partial mediation in the Coordination Game, did not hold when advantageous inequality aversion was included as an additional mediator. This may relate to the considerable reduction in sample size (N = 202 instead of N = 244) due to missing estimations of individual’s advantageous inequality aversion (see *Methods*).

### Computational modelling of updating expectations

To understand how children, adolescents, and young adults update their expectations in different social environments, we developed computational models that extend basic reinforcement learning models ^42^. In our models, participants use the outcome of interactions to update their expectations of their interaction partners’ choices in each social environment (Figure 4a-c). The extent to which these expectations are updated is reflected in a learning rate (*λ*). Besides quantifying the updating of expectations, this computational approach allows us to confirm the role of social preferences as observed in our behavioural analyses. We extended the basic reinforcement model by i) incorporating mean cohort-level social preferences to calculate a subjective value of interaction monetary outcomes (Figure 4a, 4c) that drives decision making, and ii) by allowing learning rates (expectation updating) to exponentially decay over trials of the game. Thus, we fitted four variants of this model (with and without social preferences; with and without decaying learning rates) to our experimental data for each age cohort and each game, to allow estimating different parameters (learning rates, expectation updating) across cohorts per game (see *Methods*).

**Figure 4.**
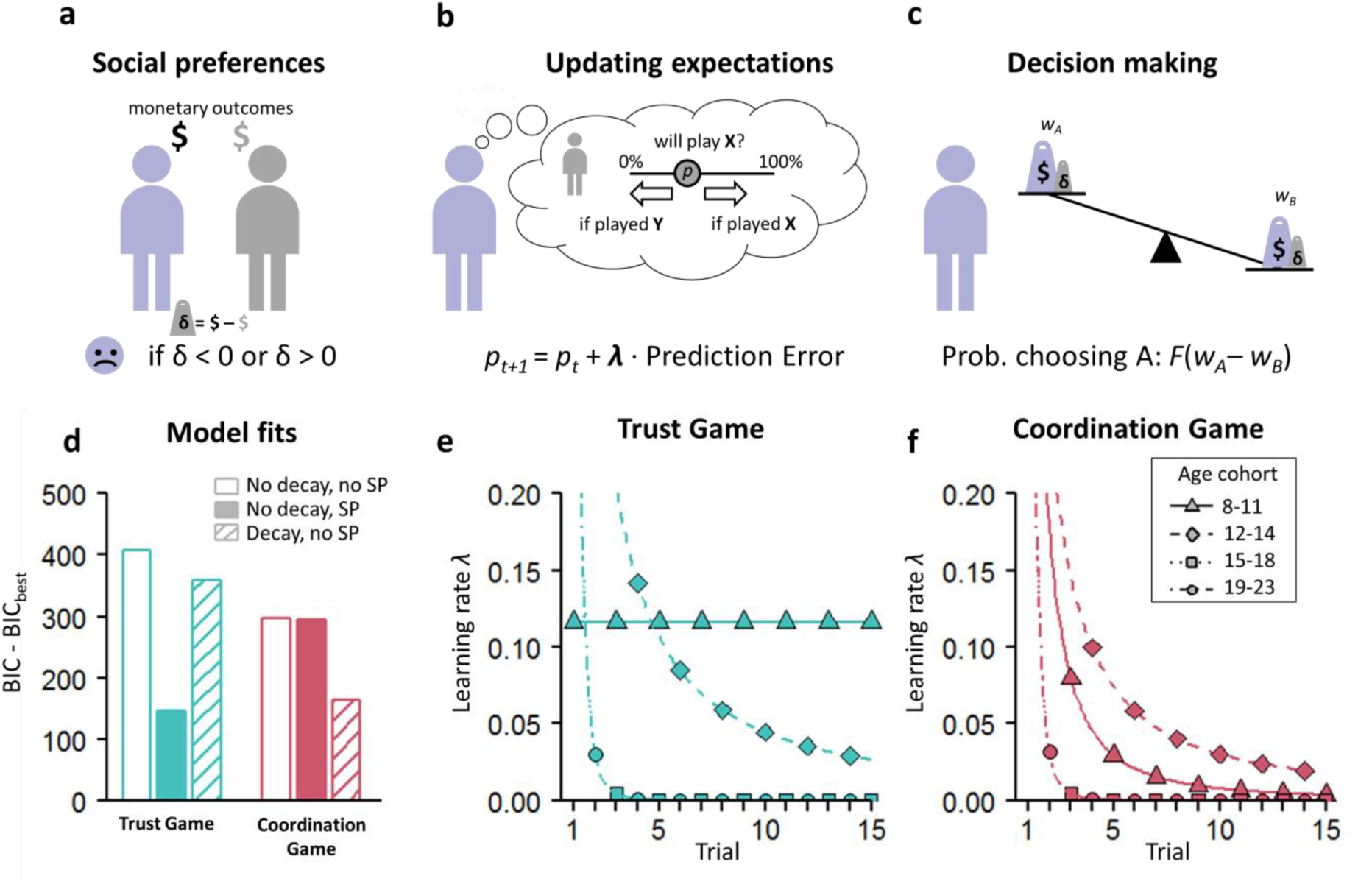
Model of reinforcement learning with social preferences. **a**, Individuals acquire monetary payoffs ($) from interactions in the games (*cf*. Figure 1). How they subjectively value each outcome can be influenced by their social preferences (Fehr & Schmidt, 1999), whose impact is proportional to the differences in payoff between interaction partners (δ). **b**, Over the course of the games, individuals update their expectations (*p*) about the behaviour of the other players. The extent of updating is proportional to the prediction error (i.e., the difference between the expected and actual outcome) and learning rate *λ* ^43^. **c**, The probability that an individual chooses A (not B) is an increasing function (*F*) of the difference in weight of the two choice options (*w*_*A*_ and *w*_*B*_; Methods). We use maximum likelihood methods to estimate learning rates *λ* for each age cohort and each game separately. In the baseline model, the weights reflect individuals’ monetary payoff from their interactions ($) and *λ* is constant over trials. Extended models also incorporate social preferences and learning rates that decay exponentially over game trials (Methods). **d**, For both games, bars show BIC differences of three model variants with the best model, which includes both social preferences (SP) and decaying learning rates. For each model, BIC values were calculated by summing BIC values of all age cohorts. Solid bars reflect models including social preferences; void and hatched bars respectively show models excluding and including decaying learning rates. **e**, Estimated learning rates from the best models as a function of the trial number, for each of the four age cohorts separately for the Trust Game, and **f**, for the Coordination Game. For the older cohorts (15-18 and 19-23), the estimated learning parameters were virtually identical, causing the plotted lines to coincide.

For both the Trust Game and the Coordination Game, a comparison of model fits provided strong support for models extended with social preferences (Figure 4d), confirming the results from our behavioural analyses that social preferences impact decision making. The best models also included decaying learning rates (see Figure 4d and Table S7). For the Trust Game, we observe that for the 8-11 year-olds, estimated learning rates are constant over the course of the game, suggesting that in late phases, individuals in this youngest age cohort still updated their expectations of the behaviour in the different social environments. In the older age cohorts, learning rates start high (around *λ*=1; asymptote not shown in Figure 4e) and decay over trials, indicating that expectations take form relatively early in the game, and remain relatively stable later on. For the Coordination Game (Figure 4f), we observe a similar pattern: older participants tended to show the strongest decay in learning rates over trials, whereas participants from the younger cohorts tended to update their expectations more early in the game.

## Discussion

Here, we examined children’s, adolescents’ and adults’ ability to learn to adjust to social environments that differ in their level of cooperation, and examined the role of social preferences (inequality aversion), prior expectations about others’ behaviour, and the updating of expectations as potential mechanisms in this behaviour. To this end, participants played a series of economic games with groups of age-matched unfamiliar others, which captured two important cooperative behaviours: trust and coordination behaviour. Our results show a striking developmental asymmetry in the learning to adjust (un)cooperative behaviour: people adjust well to environments that require uncooperative behaviour (i.e., withholding trust, putting oneself ahead) from a young age, yet only learn to adjust to environments that require cooperative behaviours during adolescence. Thus, the chances that cooperative interactions emerge differ substantially between developmental windows. Our results provide several insights into the mechanisms that explain these age-related differences.

First, age-related differences in learning to adjust to cooperative behaviours can be partly explained by differences in social preferences. Specifically, older participants showed lower levels of disadvantageous inequality aversion which explained their higher levels of cooperative behaviours. That is, younger participants are less willing to cooperate given that they are more averse to potential non-cooperation of the other player. Moreover, our computational models confirmed that participants’ decisions were best captured by a reinforcement learning model extended with social preferences. Together, these results underline that for understanding age-related changes in social - decision making it is critical to understand the development in social preferences, which differ across developmental windows and largely drive social decision-making.

A potential mechanism that may relate to the influence of inequality aversion on decision making, is behavioural control ^22^. Behavioural control refers to the ability to control thoughts and actions in order to regulate behaviour towards (long-term) goals ^44,45^. Developmental studies have shown that behavioural control undergoes protracted development due to a prolonged maturation of underlying neural circuitry in regulatory brain regions including the prefrontal cortex ^45–47^. In turn this would result in developmental changes in responses to inequality into childhood and presumably into adolescence ^23,48^. An experiment in children also confirmed a direct role of behavioural control in behaviour that benefits others: taxing children with a response inhibition task resulted in less prosocial behaviour and more costly punishment to violations of fairness ^49^. An alternative explanation is that inequality may evoke stronger emotional responses, such as increased levels of anger ^19,50^. This would yield a different view on social preferences in which responses to inequality can be based on emotion regulation ability. Future studies may further disentangle the role of behavioural control and emotion regulation as self-regulatory processes that may drive the development of social preferences and cooperative behaviours. Our findings are consistent with the idea that self-regulatory processes may be a mechanism that attunes cooperative (coordination with unequal outcomes and trust) behaviours more broadly. Consequently, an interesting field for future studies is whether strengthening self-regulatory processes is a promising pathway for stimulating cooperative behaviour in young people.

Besides social preferences, we also examined how people’s *prior expectations* of others’ trustworthiness and inclination to take more than others influenced learning in different social environments. Our results indicated that reported prior expectations of others’ behaviour were stable across age cohorts. This is surprising given the consistently reported increase in cooperativeness across age (e.g., ^5,12–14^), which was also observed in the current experiment. This suggests that there is a developmental mismatch between prior expectations and the actual levels of cooperation. Moreover, contrary to our hypotheses, we did not find effects of prior expectations on learning to adjust to different social environments. Perhaps people do not have strong prior expectations about others’ behaviour in the anonymous games used in the current study, and any expectations they might have are overridden quickly by outcomes of interactions. Presumably, effects of prior expectations in the current setup would be more prominent in a more heterogeneous sample with greater diversity in - for example - life-history backgrounds. For instance, prior expectations (as well as the updating of these expectations), may be different for people who have grown up in an environment where rewards and punishments are unpredictable, this may be particularly the case for children who have experienced harsh and inconsistent discipline, maltreatment and neglect ^51,52^. These expectations of others’ behaviour may match their environmental experiences and as such, they may engage in social situations differently. Thus, when assessing the generalizability of our results it would be important to include a more heterogeneous sample with greater diversity in life-history backgrounds. Including different populations could also help answering the question to what extent prior expectations about behaviour in games reflect prior expectations about cooperative behaviour in the real world (e.g., ^53^).

Here we used computational modelling to quantify how quickly children and adolescents updated their expectations based on choice outcomes in previous interactions. Interestingly, when placed in a new social environment, people were initially highly sensitive to behaviours of other players, and quickly adapted their behaviour to the outcomes they experienced. For older ages, behaviour stabilized after a few interactions as signalled by a decrease in learning rate. Children and young-adolescents, however, continued to react to the choices of others across the games. That is, they often switched strategies after a surprising response from one of the environments. This finding indicates that during adolescence, people more effectively integrate outcomes over time, and consequently form stable expectations of others based on their behaviour, which are not quickly overridden by a single experience. Building lasting relations may crucially depend on this integrated information of others’ behaviour. Although the continuous expectation updating of children and adolescents hampers their learning in stable environments, this actually may provide an advantage in fast-changing or unpredictable environments ^54^. That is, in such environments, immediately responding to changing feedback is more beneficial than sticking to prior expectations ^32^. Whether fast-updating better fits children’s and adolescents’ experienced social environments is an interesting question for future studies.

In the current study, participants were confronted with choices from actual peers and real-life consequences of their actions for all interaction partners. This two-directional approach, rather than often-used one-way decision making, is acknowledged as an important aspect of paradigms in social sciences ^55^. However, the controlled social environments in our study are less complex than real-life social interactions, in which factors such as social status, culture, or reputation may complicate social decision making. Future studies, e.g., field studies or studies using virtual reality, could aim to further approach the complexity of real-life social interactions, while retaining experimental control. In addition, we included a specific experimental set-up of social learning in which participants were given prior information on the different social environments. Future studies will need to assess whether our developmental findings hold in settings where participants need to figure out base rates of cooperativeness and exploitation on their own. Another limitation of the current study is that whereas social preferences were revealed preferences, prior expectations were stated expectations about others. People find it hard to estimate probabilities, and future studies need to assess the validity of these preferences with individual difference measures. Moreover, although IQ did not differ between groups and did not influence any of our findings, our adult participants were mainly recruited through university advertisements. Future studies should aim for a representative sampling strategy in each age cohort. A final limitation of the current study is its cross-sectional design, as longitudinal studies are necessary to identify developmental patterns. Therefore, developmental interpretations of behavioural results and the underlying mechanisms remain speculative.

In sum, we combined computational learning models and experimental social manipulations to demonstrate age-related changes in adjusting cooperative behaviours. Well-developed social skills are essential for succeeding in society and long-term positive outcomes. The ability to adapt to different social environments and discern who we should trust and cooperate with, may benefit short-term outcomes, but may also foster social relationships and restrain behavioural and mental health problems in the long-term ^1–4^. Knowledge of how such social skills manifest in different developmental stages inform what ages are the important developmental phase for monitoring social development, and what ages are potentially more receptive to interventions ^2,56^.

Our study has shown that adjusting cooperative behaviours is developing rapidly in early adolescence. Improvements in adjustment to different social environments are driven by developing social preferences (waning aversion to disadvantageous inequality aversion) and increasingly effective updating of own behaviour in response to others’ behaviour. Early adolescence would, therefore, be a key target window for interventions targeted at stimulating cooperative and well-adjusted social behaviour. Moreover, these findings provide important starting points for interventions for youth with maladaptive social tendencies, such as youth with conduct disorder problems ^57,58^.

## Methods

### Participants

A total of 269 participants (58.4% female) between ages 8 and 23 years took part in this study. Participants were recruited from a primary school (*n* = 60), two secondary schools (*n* = 128), and through local advertisements at a university campus (*n* = 81) in the western and middle part of The Netherlands. The majority of the participants (92.3%) were born in the Netherlands, and a minority was born elsewhere (Marocco 1.4%; all other countries <1%), or information was missing (1.4%). Twenty participants from secondary schools (ages 14-16) were excluded due to technical problems with saving the learning data. Four participants were excluded because they did not finish the cognitive behavioural measures, and therefore IQ could not be estimated. The final sample consisted of 245 individuals aged between 8 and 23 years.

Adult participants provided written informed consent. For minors, written informed consent was obtained from parents. To make the tasks incentive-compatible, participants were informed that with each behavioural task they could win points that represented lottery tickets. In each class, and in a similar-size group of adults, one lottery ticket was randomly drawn and the winner received a digital 10 Euro gift voucher. In addition, all minors received a small gift; adults received 10 Euros flat rate or course credit. All procedures were approved by the Psychology Research Ethics Committee of Leiden University (minors: CEP17-0301/120; young adults: CEP17-1009/334) and performed in accordance with the relevant guidelines and regulations.

For analyses using age cohorts (see *Computational modelling* in the section below), we divided the sample into four roughly equally-sized age cohorts: 8-11 year-olds (n=54, 46.3% female, mean age 10.6, SD 0.9), 12-14 year-olds (n=73, 52.1% female, mean age 13.4, SD 0.7), 15-18 year-olds (n=57, 59.6% female, mean age 17.0, SD 1.3), and 19-23 year-olds (n=61, 80.3% female, mean age 21.1, SD 1.4). A *χ*^2^-test indicated sex differences between age cohorts 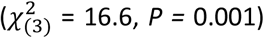, with more females in the oldest age cohort. IQ was estimated using a speeded version of the Raven Standard Progressive Matrices^59^. The estimated IQ scores were largely within the normal range varying between 79 and 136 (mean IQ = 106, SD = 10.3), and did not differ significantly between age cohorts (*F*(3,237) = 2.18, *P* = 0.090) and sexes (*F*(1,237) = 0.28, *P* = 0.770). Additional analyses showed that sex differences and IQ did not confound performance on the social games, and did not influence any of our observed age-related changes therein (see Tables S2 and S4).

### Pre-test

A key component of the economic games used in the current study is that choices have consequences not only for oneself, but also for the other player. To ensure this, we performed a pre-test at a separate high school and a separate adult sample (both in The Netherlands) functioning primarily as a match for determining the participants’ outcomes and thereby creating a true social consequence of behaviour.

In total, 82 adolescents and 44 adults were asked to make one choice (X or Y) for each social game (Trust Game and Coordination Game). We randomly linked each participant in the full-experiment with one pre-test participant. This match and the combined outcomes of their choices determined the outcome for the participants (number of points), as well as for the pre-test participant. The pre-test participants had a similar lottery ticket procedure as the participants from the full experiment, i.e., points were lottery tickets with which they had a chance of winning a 10 Euro gift voucher. All pre-test participants received a similar instruction as the participants of the main study. That is, it was stressed that their choices would have consequences for themselves and another participant, since their outcomes would result from their combined choices.

### Economic games: Trust Game and Coordination Game

Participants completed two incentivized economic games: A Trust Game and a Coordination Game (Figure 1). Each game was composed of 30 trials in total: each trial was a one-shot game with a new anonymous player (whose decision had been recorded in the pre-test; see above). Every trial the participants chose between 2 options (A or B) to distribute points between themselves and the other. After their decision they could see the choice of the player (X or Y) and the outcomes for themselves and the player. Outcomes for self and the player resulted from their combined choices, as shown with payoff matrix 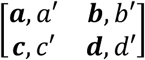 where in each of the cells entries with and without apostrophes indicate payoffs for, respectively, the other and self (in bold).

In each of the games, the two social environments consisted of 20 players each (but note that participants interacted with only 15 players per environment). Environments are formed based on pre-test responses, which were matched to create a ‘cooperative’ (73%, i.e., 11 out of 15) and an ‘uncooperative’ social environment (Figure 1). Over the course of the game trials, participants could learn the tendency of choosing X for each environment of other players, and adjust their responses accordingly. Participants were incentivised by associating their performance to the chance of winning a gift voucher (see Supplementary Information for the instruction protocol).

The Trust Game (Figure 1b) was characterized by payoff matrix 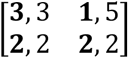. Participants could maximise their earnings by choosing A (‘trust’; top row) when matched with a member of the trustworthy environment, and choosing B (‘not-trust’; bottom row) when matched with a member of the untrustworthy environment. The Coordination Game (Figure 1d) was characterized by payoff matrix 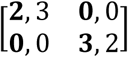. Participants could maximize their earnings by matching their partners’ choices, but the one equilibrium (A, X) put the partner ahead, while the other equilibrium put the participant ahead (B, Y).

The order of these two games was counterbalanced across participants. Within each game, participants played 30 trials, 15 trials with each environment of players (e.g., trustworthy and untrustworthy environment). The inconsistent choices within an environment (e.g., Y when playing with someone of the environment hat prefers X) were distributed across trials, yet fixed on trials 4, 8, 12, and 14. Within a game, the order of interactions with the two different environments was presented randomly, yet fixed across participants.

Although our main research questions center on the factors specific to learning to adjust behaviour in different social environments (e.g., the role of prior expectations about others, and getting more or less than others), we also included a non-social learning task to examine the level of behavioural adjustment in a simple learning context (Figure S1 and Tables S5-S6). In this non-social learning task, participants played with computers as interaction partners, and only the participant – not the computers – could receive payoffs. A formal comparison between age-related changes in learning to adjust to non-social versus social environments is included in the Supplementary Information. A computational modelling approach on the non-social game is discussed in Figure S4.

### Social preferences

We measured disadvantageous inequality aversion and advantageous inequality aversion in two separate tasks: respectively, a modified Dictator Game (DG) and Ultimatum Game (UG). These measures were derived from an adapted (i.e., child-friendly and short) version of a DG and UG (based on ^60,61^). Participants always performed the DG and UG right before the economic games.

In the Dictator Game participants were given six binary choices to divide 10 points between themselves and another anonymous participant in the study; one option was always an unequal distribution (10/0; 10 points for self, 0 points for the recipient) and the other option an equal distribution of points for themselves and the recipient (i.e., starting with (5, 5) and decreasing to (0, 0) with each subsequent trial; (4, 4), (3, 3), (2, 2), (1, 1), (0, 0).

In the Ultimatum Game, participants responded to six proposals of another anonymous participant in the study on how to divide 10 points. In the case of a rejection both players earn zero, whereas if the participant accepted the offer, the players get the proposed outcome. The first proposal was an equal split but every next proposal was more beneficial for the other than for self (i.e., (5, 5), (4, 6), (3, 7), (2, 8), (1, 9), (0, 10). For both games, we were interested in the point at which a participant switched their preference from an equal to unequal distribution, or vice versa. This allowed us to infer the point at which participants were indifferent between either distribution. This ‘indifference point’ was used to calculate their inequality aversion (^20^; see Supplemental Information).

The values for individuals’ disadvantageous inequality aversion varied in discrete logarithmic steps between 0 and 4.5, and values of advantageous inequality aversion varied in linear steps between 0 and 1 (see, e.g. ^60^). To ensure that the non-linearity of the UG values did not influence our results, we repeated our GLMMs with participants’ calculated indifference points. Note that all of our findings remain the same when using indifference points instead of estimated social preferences in analyses.

Finally, our estimates of participants’ social preferences depend on their level of consistency in choice behaviour in the DG and UG. In total, 54 participants were excluded due to missing values for social preferences (missing disadvantageous inequality aversion, *n* = 1; missing advantageous inequality aversion, *n* = 53). See Supplementary Information, and Figure S2 and S3 for a more detailed description of the Dictator game and Ultimatum Game, and calculation of inequality aversion measures.

### Prior expectations

Before the start of each of the economic games (Trust Game and Coordination Game), we assessed participants’ prior expectations about the behaviour of other people. We asked participants “Suppose that there are 10 other players, how many of these 10 do you think will choose X?” (i.e., ‘trustworthy’ choice in the Trust Game, or ‘other-ahead’ choice in the Coordination Game). This resulted in a prior expectation of the trustworthiness of others (Figure 3c) and tendencies to get more than others (Figure 3d), varying from 0 to 10.

### Procedure

All tests were administered in school settings. In the instruction of each learning task, three control questions were included to ensure understanding of the experimental procedure. Two questions quizzed the participant on their understanding of the point distribution (e.g., type how many points each player was winning in a certain choice combination), and one question referred to the colour denotation of the two environments. If participants failed one of the control questions, the instruction was repeated until participants understood the procedure of the game. For participants younger than 12, instructions were read out load by an experimenter. All participants completed the tasks by themselves on computers in a quiet environment at school or at the university. Background variables such as the Raven SPM (estimated IQ) and several questionnaires (not relevant to the current study) were administered online using Qualtrics (www.qualtrics.com). In a separate session the DG, UG, and learning tasks were completed using the online software LIONESS Lab ^62^

### Statistical analyses of behavioural data

To assess age-related changes in prior expectations and social preferences we ran separate robust linear regression analyses (5000 bootstraps), each with age linear and age quadratic as predictors. Multiple mediation analyses were conducted in SPSS using the computational tool PROCESS version 3.3^63^. For indirect effects, 95% (two-tailed) bias-corrected bootstrapped confidence intervals were calculated using 5000 repetitions. An indirect effect is significant if the confidence interval for the indirect effect does not include zero. These analyses were conducted in SPSS 25, and all tests were two-sided.

### Generalized linear mixed models

To analyse choice behaviour in the Trust Game and Coordination Game, we fitted logistic generalized linear mixed models (GLMMs) to decisions to choose A (coded as 0) or B (coded as 1) to each game separately. Analyses were conducted in R 3.6.1 ^64^, using the lme4 package ^65^). In all models, participant ID entered the regression as a random intercept to handle the repeated nature of the data. Where appropriate, environment was entered as a random slope in our analyses to handle the differences between individuals in their responsiveness to learning to different levels of (non)cooperation. Our GLMMs included a main effect of environment (e.g., trustworthy environment, untrustworthy environment), age in years (linear and quadratic), prior expectations of others’ choices and social preferences, and all two-way interactions with environment (see Tables S1-S4 for all GLMM results). Note that for the Trust Game we only added disadvantageous inequality aversion, whereas for the Coordination Game both social preferences were included. That is, in the Coordination Game both types of inequality can occur and drive choice behaviour, in contrast to the Trust Game in which only disadvantageous inequality is present.

In all GLMMs, age, prior expectations, disadvantageous and advantageous inequality aversion (mean-centered and scaled) and categorical predictor variables were specified by a sum-to-zero contrast (e.g., sex: -1 = boy, 1 = girl). For the mixed-effects model analyses the optimizer “bobyqa” ^67^ was used, with a maximum number of 1×10^5^ iterations. P-values for all individual terms were determined by Loglikelihood Ratio Tests as implemented in the mixed function in the afex package^68^. All statistics, including odds ratios and confidence intervals, are reported in Tables S1 –S 6.

### Computational modelling

To gain a mechanistic understanding of participants’ learning to adjust in the Trust Game and the Coordination Game, we used a basic reinforcement learning (RL) model^43^ and extended it to accommodate social preferences ^20^ (aversion to unequal outcomes). All our models follow the basic logic of RL, in which agents learn about others behaviour by updating their expectations with experience. In the case of the games, these expectations (denoted *p*) concern the behaviour of their interaction partners (X or Y; cf. Figure 1). In each trial, *p* is updated with a magnitude proportional to the prediction error (PE; the difference between the actual and expected choice) and the learning rate *λ*. Formally, *p*_*t+1*_ = *p*_*t*_ + *λ* · *PE*, where *PE* = *p* – choice of other (1 if *X*, 0 otherwise). We fit a set of reinforcement learning models to the data to investigate how *λ* changes across age cohorts. This parameter is bounded between 0 (which means no updating of expectations at all) and 1 (which means that expectations match the decision of the most recent player).

In our models, the value of *p* determines the relative weights of *w*_*A*_ and *w*_*B*_ (Figure 4). Each of the games is characterized by a payoff matrix (Figure 1). In each trial *t*, expected monetary payoffs of choosing *A* or *B* are respectively given by *w*_*A,t*_= *p*_*t*_ · *a* + (1 – *p*_*t*_) · *b* and *w*_*B,t*_= *p*_*t*_ · *c* + (1 – *p*_*t*_) · *d*. We set the initial value of *p*_*0*_ to the cohort mean prior measured in our experiment. The probability that a participant chooses *A* is determined by a standard softmax function: 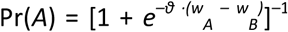. As there are only two options (A and B) to choose from, the probability of choosing *B* is simply 1 – Pr(*A*). In the softmax formula, *ϑ* reflects ‘decision sensitivity’ and accounts for stochasticity in participants’ choices: low values of *ϑ* indicate high levels of stochasticity (Pr(*A*) and Pr(*B*) tend to be near 0.5), and high values of *ϑ* indicate low levels of stochasticity. In our model fits, *ϑ* is a free parameter allowed to vary between 0 and 5.

We extended this baseline model with two factors. First, we include the cohort mean measures of social preferences; that is, we add the measured cohort averages of *disadvantageous* and *advantageous inequality aversion* to calculate of *w*_*A*_ and *w*_*B*._. In particular, for the Trust Game, the weight of option *A* was penalized with a value proportional to the disadvantageous inequality aversion (i.e., *α*; note that we drop the subscripts as we assume social preferences to be parameters with a constant value^20^ : *w*_*A*_= *p* · *a* + (1 – *p*_*t*_) · [*b* – *α* · (*b’*– *b*)]. As for option *B* the payoffs for both partners are always equal, *w*_*B*_ is unaffected by social preferences. For the Coordination Game, social preferences can affect the weights of both *A* and *B*: *w*_*A*_= *p* · *α* · (*a’*– *a*), and *w*_*B*_= *p* · *β* · (*d – d’*), where *β* denotes advantageous inequality aversion.

Second, we allowed the learning rate *λ* to decay over the course of interactions. We implemented this by defining *λ*_*t*_=*λ*_*0*_ · *r* ^-*τ*^, where *r* denotes the trial number, and *τ* is a free parameter that reflects the speed of the decay in learning, allowed to vary between 0 and 5. The values of the estimated parameters (*ϑ, λ, τ*) per age cohort and per game can be found in Table S7. For each of the four age cohorts from Figure 2 separately, we pooled the data and fitted the model with each possible combination of the factors ‘social preferences’ and ‘decay’, yielding a total of four models per cohort per game. Note that we also evaluated a potential role for prior expectations by including mean cohort-level prior expectations in the initial valuation of the choice options. However, because prior expectations were relatively close to 5 (range 0 – 10; Figure 2) this was close to the default expectation *p* of 0.5, marking indifference between the environments at the first choice. Hence, we did not apply formal tests of improved model fit for prior expectations.

Figure 4d shows the goodness-of-fit for each model summed across the four age cohorts relative to the best model, which includes both social preferences and decay. We included a simulation study with a parameter recovery component in the Supplementary Information. Our approach of fitting reinforcement learning models to cohort-level data was motivated by the fact that we did have a limited number of observations to accurately fit our model to individual-level choice data. Note that sensitivity analyses with individually-derived parameters indicated this did not influence any of our model-fit conclusions or main findings.

### Data availability

The data that support the findings of this study will be made available in the Leiden repository: https://openaccess.leidenuniv.nl.

### Code availability

All relevant R codes will be made available in the Leiden repository: https://openaccess.leidenuniv.nl.

## Acknowledgements

We would like to thank all participants and their parents, and the participating schools for their cooperation. We thank Joyce van Amstel, Melanie van Berkel, Annemarijn de Bruin, Yolinda Davidse, Ellina Guijt, Tim Habermehl, Tosca Hunink, Maaike Jacobs, Behazin Khosravi, Sophie van de Leur, Deveney Kok-Sey-Tjong, Liesbeth Roerade, Tess van der Toorn, and Iris Willink for their help with data collection. We thank Jungsun Yoo for help with programming the task, Ruth Roberts for help with formulating the child friendly task instructions, and Eveline Crone for helpful discussions. This work was supported by an Open Research Area (ORA) grant [grant number 464-15-176] financed by the Netherlands Organization for Scientific Research (NWO), the German Research Foundation (DFG) and the Economic and Social Research Council (ESRC). The funder had no role in the conceptualization, design, data collection, analysis, decision to publish, or preparation of the manuscript.

## Author contributions

B.W., L.M., W.B., E.V., and A.C.K.D. designed the experiment; L.M. programmed the economic games; B.W. collected the data; B.W. and A.C.K.D. performed the behavioural analyses; L.M., W.B., and A.C.K.D. performed the computational modelling; B.W., A.C.K.D, and L.M wrote the main manuscript text; B.W. prepared figures 1-3; L.M. prepared figure 4; all authors interpreted the results and reviewed the manuscript.

