## Supplementary Information for "Developmental asymmetries in learning to adjust to cooperative and uncooperative environments"

**This file includes:**

Figures S1 to S5

Tables S1 to S7

Supplementary Information and results

References for Supplementary Information reference citations

### Supplementary Information

#### 1. Prior expectations and social preferences: IQ, and sex effects

In a series of robust regression analyses (5000 bootstraps) we examined the effect of IQ and sex effects on participant's prior expectations and social preferences across all ages. We found that estimated IQ was not related to advantageous inequality aversion, prior expectations of other's trustworthiness, and prior expectations of other's tendency to choose more for oneself (all *P*s > 0.1). However, participants with higher IQ were less disadvantageous inequality averse (IQ, B = -0.440, β = -0.284, *P* < 0.001, 95% CI [-0.062, -0.024]). Also, compared to boys, girls scored lower on disadvantageous inequality aversion (Sex, B = -0.702, β = -0.217, *P* = 0.001, 95% CI [-1.190, -0.286]) and higher on advantageous inequality aversion (Sex, B = 1.69, β = .252, *P* = .001, 95% CI [0.730, 0.256]). Compared to boys, girls also expected that others would choose to have more than the participant (sex, B = 0.437, β = 0.129, *P* = 0.048, 95% CI [0.000 - 0.863]). We observed no sex differences in prior expectations of other's trust behaviour (all *P*s > 0.05).

Next, we assessed correlations between advantageous and disadvantageous inequality aversion, and between the prior expectations of the Trust Game and Coordination Game. Inequality aversions were slightly negatively correlated, indicating that people who disliked being behind also liked being ahead (*r* = -0.181, *P* = 0.010; N = 202). This relation was significant even when controlled for age (*r* = -0.151, *P* = 0.032). Prior expectations about others in the Trust Game and the Coordination Game were not correlated (*r* = 0.069, *P* = 0.282; N = 245), neither when controlled for age (*r* = 0.067, *P* = 0.297).

#### 2. Developmental changes in the non-social learning task

To assess whether people are able to adjust their decision-making in a non-social context, we included a non-social learning task with two environments of computer opponents (payoff matrix $\begin{bmatrix} \mathbf{3} & \mathbf{0} \\ \mathbf{0} & \mathbf{3} \end{bmatrix}$, see Figure S1a). Similar to the economic games, participants could maximise their payoffs by coordinating their choices to the choices of the computer opponent. That is, choosing option A when playing against the environment that most often (11/15 trials) chooses X, and choosing option B when playing against the environment that most often (11/15 trials) chooses Y.

We fitted a logistic GLMM to decisions (coded as correct/incorrect) in the non-social learning task, with age linear and age quadratic as predictors. Performance increased linearly with age (Figure S1b; see Table S5 for all statistics).

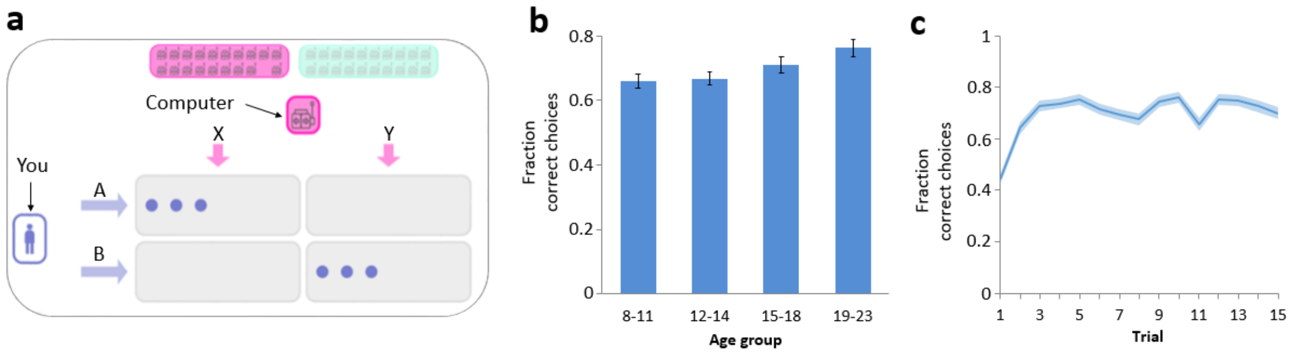


**Figure S1. Non-social learning task.**
**a**, layout of the learning task. **b**, performance per age cohort. **c**, performance over trials pooled across all participants (N=245). Error bars in panel b and shaded areas in panel c show standard error of the mean (s.e.m).

#### 3. Age-related differences in the non-social vs. social learning game

The main focus of this study was to compare the influence of prior beliefs, social preferences, and feedback-based updating across age groups in different social learning environments. However, the inclusion of a non-social task lends itself for the opportunity to compare learning performance in a social and non-social task more generally. Exploratively, we therefore compared performance in the non-social learning task versus the Trust Game and the non-social task versus the Coordination Game.

We fitted a binomial GLMM to decisions (coded as correct/incorrect) with age linear * type of task (social vs non-social) as predictors. Participant was included as a random intercept to account for the repeated nature of the choice data. Overall performance (grouped across environments) was lower in the Trust Game compared to the non-social learning task; main effect Task $P$ <0.001; Trust Game *Mean* accuracy = 0.613 (*SD* = 0.487); Non-social *Mean* accuracy = 0.699 (*SD* = 0.459). Follow-up tests, showed that this advantage for learning in the non-social task was present for each age group (8-11 years $P$ <0.001; 12-14 years $P$ <0.001; 15-18 years $P$ = 0.024; 19-23 years $P$ <0.001). A similar GLMM showed that overall performance (grouped across environments) was also lower in the Coordination Game compared to the non-social task; main effect Task $P$ <0.001; Coordination Game *Mean* accuracy = 0.668 (*SD* = 0.470). Follow-up tests showed that this was, however, only the case for the two youngest age groups, and from approximately age 15 adolescents performed -on average- equally well in both games (8-11 years $P$ < 0.001; 12-14 years $P$ < 0.001; 15-18 years $P$ = 0.870; 19-23 years $P$ = 0.830). These comparisons between games confirm that specifically young adolescents find it difficult to adjust behaviour in social compared to non-social games. Our analyses described in the main paper shed light on what factors underlie age-related change in learning in social environments that differ in their level of cooperation.

#### 4. Testing confounding effects of sex and IQ on choice behaviour in the social-learning games

To assess possible effects of sex and IQ on choice behaviour in the learning tasks, we ran additional GLMMs (one per game) that included all predictor variables from the main model (see *General linear mixed-effects models* in main text), and added main effects of sex and estimated IQ. In the Trust Game there were no main effects of sex or IQ (see SI table 2 for all statistics); all effects remained significant after adding sex and IQ to the model – except for the main effect of disadvantageous inequality aversion. In the Coordination Game, there were also no main effects of sex and IQ (see SI table 4 for all statistics). All effects remained significant after adding sex and IQ to the model. In the Non-social learning task, there was no main effect of sex (see SI table 6 for all statistics). However, there was a main effect of IQ ($P < 0.001$) indicating that participants with a higher IQ had better performance in the non-social learning task. These results suggest that IQ and sex differences did not influence performance on the social learning tasks, and did not influence any of the observed age-related changes in adjustment behaviour. However, IQ did relate positively to a general learning tendency in the non-social learning task.

#### 5. Calculation of social preferences Dictator Game

The Dictator Game (DG) was used to estimate people's advantageous inequality aversion. The aim of the DG is to identify the point at which a participant is indifferent between (10, 0) and the equal distribution. To this end, we always had participants initially chose in the first trial between (10, 0) and (5, 5). The number of points in the subsequent trials depended on the choice in that first trial. That is, if in the first trial a participant selected the equal distribution (5, 5) rather than the (10, 0), the number of points in the equal distribution decreased with one point in each subsequent trial (i.e., (5, 5); (4, 4); (3, 3); (2, 2); (1, 1); (0, 0)). In contrast, if in the first trial the participant selected the unequal distribution (10, 0) over the equal distribution (5, 5), in the subsequent trials the number of points in the equal distribution increased to (10, 10) (i.e., (5, 5); (6, 6); (7, 7); (8, 8); (9, 9); (10, 10)). A participant's inequality aversion is determined by the equal distribution (x, x) which he/she regards as good as the (unequal) distribution (10, 0). For participants with consistent preferences (YY%), this 'indifference point' (*IP*) follows from the point at which they switch from choosing the equal distribution over the unequal distribution (or vice versa; Blanco et al., 2011). For example, if a participant prefers (10, 0) over (6, 6), but prefers (7, 7) over (10, 0), we assume that $IP$=6.5. Following Blanco et al., (2011), we calculate a participant's advantageous inequality aversion as: $\beta = 1 - IP/10$.

We used the general constraints as formalized by Fehr & Schmidt (1999), with the range of advantageous inequality aversion values limited between 0 and 1. For participants who switched multiple times from equal to unequal distribution (i.e., who did not show a consistent choice pattern), we fitted a softmax function (see below) to their DG choices to approximate their indifference point. In particular, we coded a choice for the unequal distribution as 0 and the equal distribution ($x$, $x$) as 1. To these choices, 6 for each participant, we then fitted a softmax function $y=[1+\exp(-Z)]^{-1}$, where $Z = a + b*x$. Subsequently, we solved for y=0.5 (i.e., $-a / b$) if the fitted value of $b$ was positive (i.e., when participants tended to choose equal outcomes more often when the value of $x$ increased). If the approximated indifference point was outside of the theoretically possible range of [0,10], we assumed it was undefined. This procedure resulted in approximated switch points *IP'*, and we calculated the participant's advantageous inequality aversion in the same way as above: $\beta=1 - IP'/10$.

In the Dictator Game, 148 participants showed a consistent choice pattern (i.e., up to 1 switching point, 60%). For 54 participants with multiple switching points we successfully used the softmax function to approximate their indifference point. For 43 participants the approximated indifference point was outside of the theoretically possible range [0,5], or had a negative values of $b$. These participants where therefore excluded from analyses that used the advantageous inequality aversion measure.

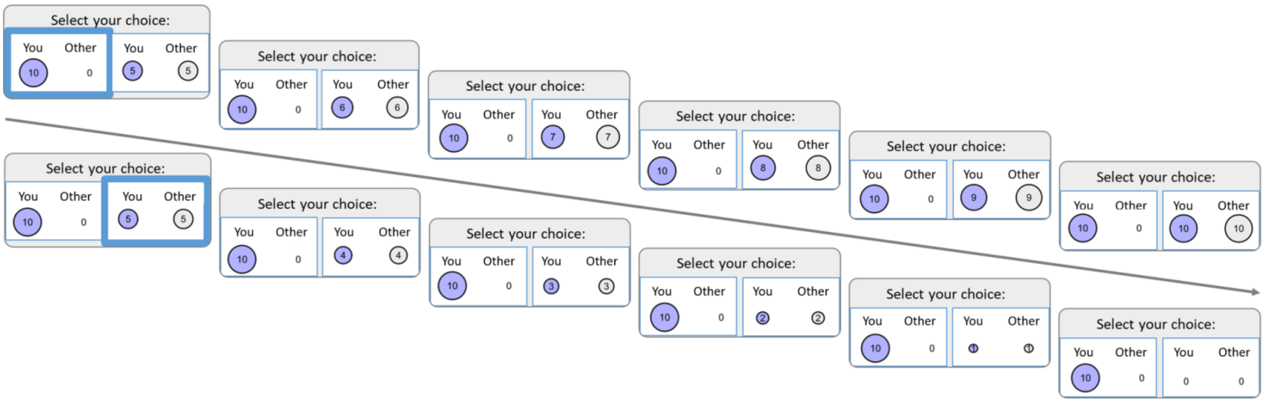


Figure S2. Display of the Dictator Game trials.
Trial sequence depends on the choice in the first trial: upper sequence when first choice is for the unequal distribution, lower sequence when first choice is for the unequal distribution.

##### 6. Calculation of social preferences Ultimatum Game

The Ultimatum Game (UG) was used to estimate people's disadvantageous inequality aversion. The UG is a sequential two-stage game, in which a proposal for a division of points is offered by the proposer, and can be rejected or accepted by the responder. In the case of a rejection both players earn zero. If the responder accepts, players get the outcome proposed. In the proposer stage participants were presented with 10 points and could make an offer to the responder keeping 0 up to 10 points for themselves (i.e., 10 (self), 0 (other); (10, 0); (9, 1); (8, 2); (7, 3); (6, 4); (5, 5); (4, 6); (3, 7); (2, 8); (1, 0)).

In the responder stage, participants were told they were now paired with a new player and that they could accept or reject their proposals (Figure S). As a responder, participants responded to 6 proposals. The first proposal was an equal split but every next proposal was more beneficial for the other than for self (i.e., (5, 5), (4, 6), (3, 7), (2, 8), (1, 9), (0, 10)). This responder stage was used to obtain a measure of disadvantageous inequality aversion, given that the minimum-acceptable offers in the UG is used as a point estimate of disadvantageous inequality aversion for each individual. We used the settings and calculations of Blanco et al., (2011) with the constraints that if there is no rejected offer, disadvantageous inequality aversion = 0 and if all offers are rejected, disadvantageous inequality aversion = 4.5. Like in the DG, we calculated participants' indifference points (*IP*). Following Blanco et al., (2011) we then calculated a participant's disadvantageous inequality aversion as: β=*IP*/(2*(5-*IP*).

In the Ultimatum Game, the majority of participants showed a consistent choice pattern (i.e., up to 1 switching point, *n* = 234, 95.5%). In total, 11 participants had multiple switching points (up to 3), and again we used the softmax function described above for the Dictator Game to approximate their indifference point. For 1 participant the approximated indifference point was outside of the theoretically possible range [0,5], or had a negative values of *b*. This participant was therefore excluded from analyses that used the disadvantageous inequality aversion measure.

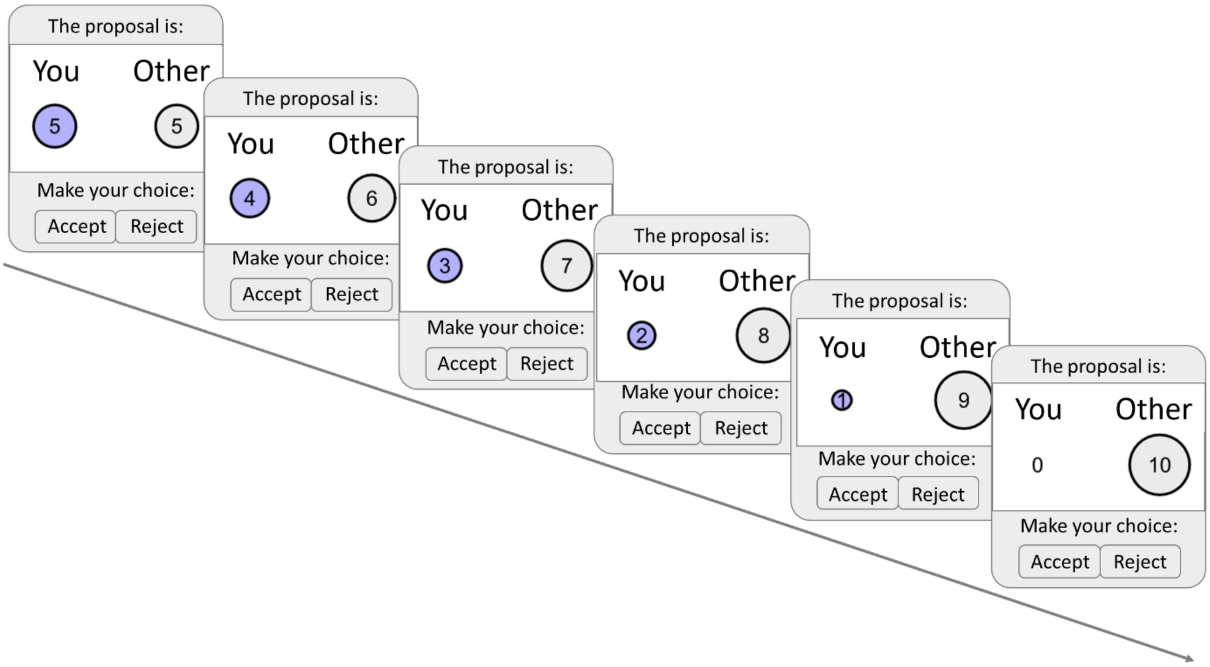


**Figure S3. Display of the Ultimatum Game trials (responder stage).**

#### 7. Computational models assessing behavioural adjustment in the non-social learning task

To assess behavioural adjustment outside of social interactions, we fitted the reinforcement models to decisions in the non-social setting (see Figure S4). Following the main text, we pool all data from participants per cohort and consider models with and without decay in learning rates across trials. As there are no other people involved, we do not consider models with or without social preferences, and assume that participants' priors in the first trial are such that they expect their computerized opponent to choose either action with equal (50%) probability.

We observe that models including a decay in learning rates lead to an improved fit. Figure S4 illustrates the estimated learning parameters of the best-fitting model. We observe that for each age cohort, learning rates decreased strongly over time. Interestingly, and in contrast to the Trust Game and the Coordination Game, the least strong decay was observed for the oldest age cohort.

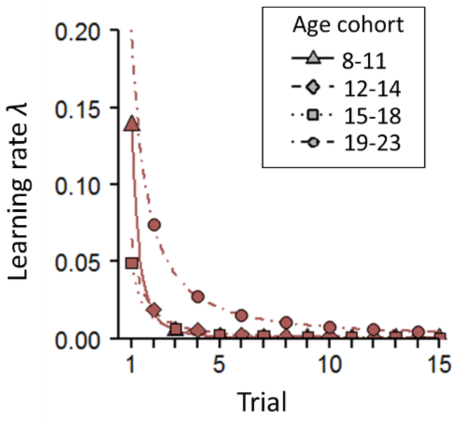


**Figure S4. Estimated learning rates per age cohort from the best models for the Non-Social learning task**

Best models include decaying learning rates. Estimates learning rates as a function of the trial number, for each of the four age cohorts separately for the Non-Social learning task.

#### 8. Recovery of computational models

To assess the robustness of the computational models presented in the main text, we performed a recovery analysis. For each of the social games (Trust Game and Coordination Game), and for each of the age cohorts separately, we used the parameters of the best fitting models (which included social preferences and a decaying learning rate) to simulate 100 mock data sets. The distribution of mock participants across cohorts was the same as in our behavioural data set. Subsequently, we fitted each of the four models (including and excluding social preferences and decaying learning rates) to the simulated data and compared their fits. This allowed us to examine the recoverability of our models. For both economic games, for each of the 100 simulated mock data sets, the model including social preferences and decaying learning rates fitted the data best, showing that our best fitting model is recoverable.

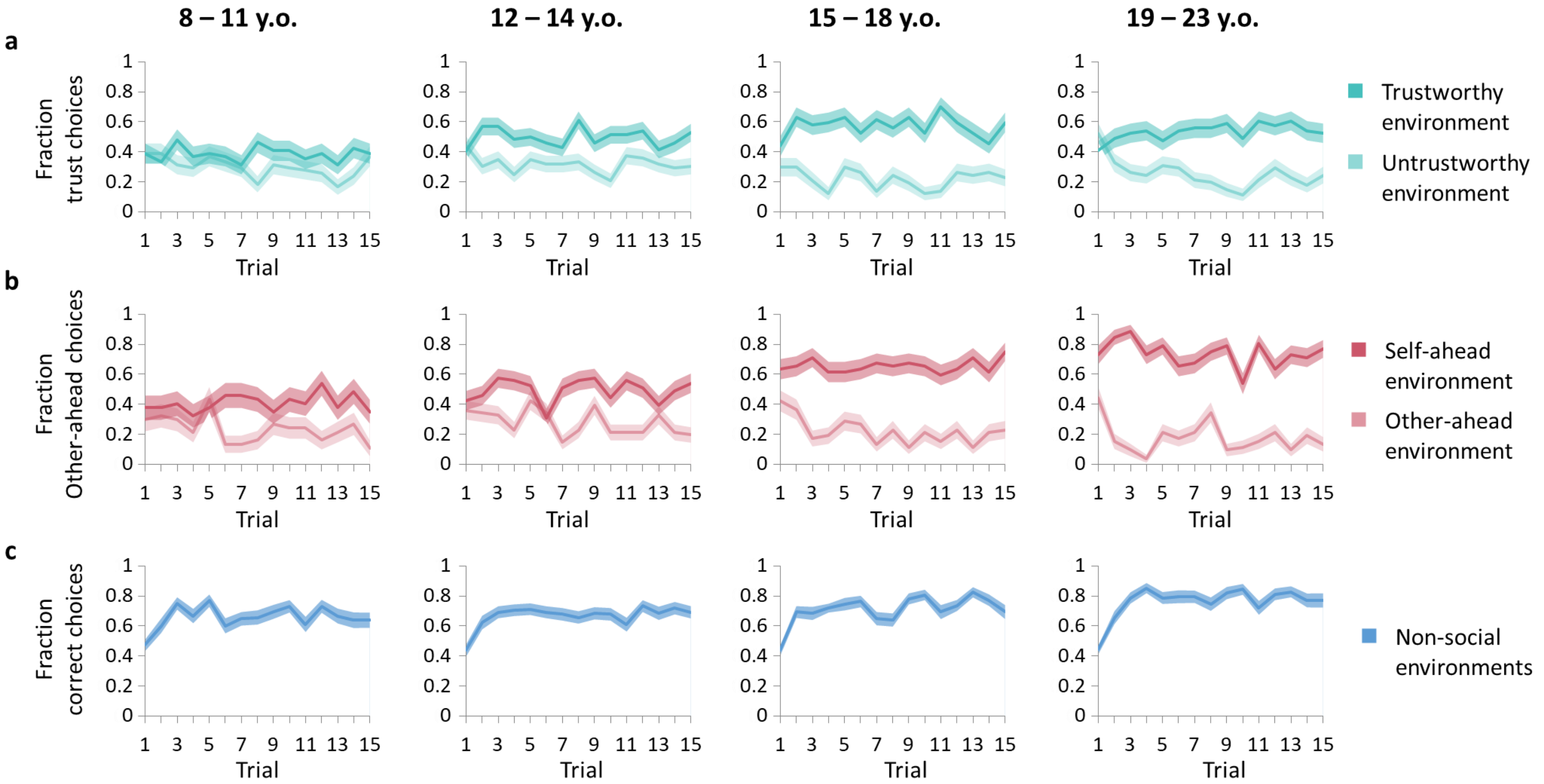


**Figure S5.**

Display of choice behaviour over trials separately for each age cohort, in (**a**) the Trust Game (N = 244), (**b**) the Coordination Game (N = 202), and (**c**) the Non-social learning task (N = 245). Shaded areas represent s.e.m.

### Supplementary Tables

#### Supplementary Table 1. Mixed-effects model for the Trust Game (N=244)

Results of the binomial generalized linear mixed model (GLMM) testing effects of age (linear and quadratic), prior expectations, and disadvantageous inequality aversion (IA), and all 2-way interactions with environment on choice behaviour in the Trust Game. Significant effects are in bold. GLMMs are described fully in the main text. Full R-code of this model is: mixed(Choice~cAge-linear*Environment + cAge-quadratic*Environment+ cDisadvantageousIA*Environment + cpriorTrustGame*Environment + (Environment | Subject), method="LRT", family=binomial, data = dat, na.action=na.exclude, control=glmerControl(optimizer = "bobyqa", optCtrl = list(maxfun = 100000)))

| | Estimate | *SE* | $\chi^2$ | *P* | *Odds Ratio* | *CI* |
|---|---|---|---|---|---|---|
| **Environment** | **-0.655** | **0.079** | **60.56** | **<.001** | **0.52** | **0.45 – 0.61** |
| Age linear | -0.074 | 0.091 | 0.637 | 0.425 | 0.93 | 0.78 – 1.11 |
| Age quadratic | 0.117 | 0.087 | 1.765 | 0.184 | 1.12 | 0.95 – 1.33 |
| **Disadvantageous IA** | **0.194** | **0.087** | **4.918** | **0.027** | **1.21** | **1.02 – 1.44** |
| Prior expectations | -0.161 | 0.083 | 3.659 | 0.056 | 0.85 | 0.72 – 1.00 |
| **Environment * Age linear** | **-0.300** | **0.087** | **11.45** | **<.001** | **0.74** | **0.62 – 0.88** |
| **Environment * Age quadratic** | **0.207** | **0.084** | **5.978** | **0.014** | **1.23** | **1.04 – 1.45** |
| Environment * Disadvantageous IA | 0.148 | 0.083 | 3.141 | 0.076 | 1.16 | 0.99 – 1.36 |
| Environment * Prior expectations | 0.107 | 0.080 | 1.783 | 0.182 | 1.11 | 0.95 – 1.30 |
| Post-hoc \| *Trustworthy environment* | | | | | | |
| **Age linear** | **-0.374** | **0.142** | **6.817** | **0.009** | **0.69** | **0.52 – 0.91** |
| **Age quadratic** | **0.322** | **0.136** | **5.569** | **0.018** | **1.38** | **1.06 – 1.80** |
| **Disadvantageous IA** | **0.341** | **0.134** | **6.313** | **0.012** | **1.41** | **1.08 – 1.83** |
| Prior expectation | -0.053 | 0.128 | 0.168 | 0.682 | 0.95 | 0.74 – 1.22 |
| Post-hoc \| *Untrustworthy environment* | | | | | | |
| **Age linear** | **0.229** | **0.109** | **4.571** | **0.037** | **1.26** | **1.01 – 1.56** |
| Age quadratic | -0.091 | 0.104 | 1.343 | 0.386 | 0.91 | 0.74 – 1.12 |
| Disadvantageous IA | 0.046 | 0.103 | 0.608 | 0.660 | 1.05 | 0.86 – 1.28 |
| **Prior expectation** | **-0.268** | **0.101** | **3.778** | **0.008** | **0.77** | **0.63 – 0.93** |

**Supplementary Table 2. Mixed-effects model with IQ and sex effects for the Trust Game (N=244)**

Results of the binomial generalized linear mixed model (GLMM) testing confounding effects of IQ and sex on choice behaviour in the Trust Game. Predictors of interest are underlined. GLMMs are described fully in the main text. Full R-code of this model is: mixed(Choice~IQ + sex + cAge-linear*Environment + cAge-quadratic*Environment+ cDisadvantageousIA*Environment + cpriorTrustGame*Environment + (Environment | Subject), method="LRT", family=binomial, data = dat, na.action=na.exclude, control=glmerControl(optimizer = "bobyqa", optCtrl = list(maxfun = 100000)))

| | Estimate | *SE* | $\chi^2$ | *P* | *Odds Ratio* | *CI* |
|---|---|---|---|---|---|---|
| <u>Sex</u> | 0.037 | 0.085 | 0.195 | 0.659 | 1.04 | 0.88 – 1.23 |
| <u>IQ</u> | 0.093 | 0.086 | 1.123 | 0.288 | 1.10 | 0.93 – 1.30 |
| Environment | -0.654 | 0.079 | 60.03 | <.001 | 0.52 | 0.45 – 0.61 |
| Age linear | -0.070 | 0.094 | 0.548 | 0.459 | 0.93 | 0.78 – 1.12 |
| Age quadratic | 0.109 | 0.088 | 1.500 | 0.221 | 1.11 | 0.94 – 1.33 |
| Disadvantageous IA | 0.213 | 0.091 | 5.386 | 0.020 | 1.24 | 1.04 – 1.48 |
| Prior expectations | -0.153 | 0.084 | 3.280 | 0.070 | 0.86 | 0.73 – 1.01 |
| Environment * Age linear | -0.300 | 0.088 | 11.443 | <.001 | 0.74 | 0.62 – 0.88 |
| Environment * Age quadratic | 0.207 | 0.084 | 5.982 | 0.014 | 1.23 | 1.04 – 1.45 |
| Environment * Disadvantageous IA | 0.149 | 0.083 | 3.153 | 0.076 | 1.16 | 0.99 – 1.37 |
| Environment * Prior expectations | 0.107 | 0.080 | 1.791 | 0.181 | 1.11 | 0.95-1.30 |

**Supplementary Table 3. Mixed-effects model for the Coordination Game (N=202)**

Results of the binomial generalized linear mixed model (GLMM) testing effects of age (linear and quadratic), prior expectations, and disadvantageous inequality aversion (IA), and all 2-way interactions with environment on choice behaviour in the Coordination Game. Significant effects are in bold. GLMMs are described fully in the main text. Full R-code of this model is: mixed(Choice~cAge-linear*Environment + cAge-quadratic*Environment+ cDisadvantageousIA*Environment + cAdvantageousIA*Environment + cpriorCoordinationGame*Environment + (Environment | Subject), method="LRT", family=binomial, data = dat, na.action=na.exclude, control=glmerControl(optimizer = "bobyqa", optCtrl = list(maxfun = 100000)))

| | Estimate | *SE* | $\chi^2$ | *P* | *Odds Ratio* | *CI* |
|---|---|---|---|---|---|---|
| **Environment** | **-1.04** | **0.086** | **115.567** | **<.001** | **0.35** | **0.30 – 0.42** |
| **Age linear** | **-0.214** | **0.071** | **8.709** | **0.003** | **0.81** | **0.70 – 0.93** |
| Age quadratic | 0.115 | 0.068 | 2.823 | 0.092 | 1.12 | 0.98 – 1.28 |
| **Disadvantageous IA** | **0.184** | **0.068** | **7.260** | **0.007** | **1.20** | **1.05 – 1.37** |
| **Advantageous IA** | **-0.162** | **0.065** | **6.038** | **0.014** | **0.85** | **0.75 – 0.97** |
| Prior expectations | 0.033 | 0.065 | 0.263 | 0.608 | 1.03 | 0.91 – 1.17 |
| **Environment * Age linear** | **-0.446** | **0.094** | **21.275** | **<.001** | **0.64** | **0.53 – 0.77** |
| Environment * Age quadratic | -0.063 | 0.090 | 0.474 | 0.491 | 1.06 | 0.89 – 1.27 |
| Environment * Disadvantageous IA | 0.091 | 0.090 | 1.002 | 0.317 | 1.09 | 0.92 – 1.31 |
| **Environment * Advantageous IA** | **-0.256** | **0.087** | **8.517** | **0.004** | **0.77** | **0.65 – 0.92** |
| Environment * Prior expectation | 0.029 | 0.086 | 0.115 | 0.734 | 1.03 | 0.87 – 1.22 |
| Post-hoc \| *Self-behind* | | | | | | |
| **Age linear** | **-0.655** | **0.131** | **23.598** | **<.001** | **0.52** | 0.40 – 0.67 |
| Age quadratic | 0.180 | 0.126 | 2.025 | 0.155 | 1.01 | 1.00 – 1.03 |
| **Disadvantageous IA** | **0.277** | **0.126** | **4.744** | **0.029** | **1.32** | 1.03 – 1.69 |
| **Advantageous IA** | **-0.410** | **0.121** | **11.038** | **<.001** | **0.66** | 0.52 – 0.84 |
| Prior expectation | 0.064 | 0.120 | 0.281 | 0.596 | 1.07 | 0.84 – 1.35 |
| Post-hoc \| *Self ahead* | | | | | | |
| **Age linear** | **0.223** | **0.102** | **4.731** | **0.030** | **1.25** | 1.02 – 1.53 |
| Age quadratic | 0.054 | 0.097 | 0.298 | 0.584 | 1.00 | 0.99 – 1.02 |
| Disadvantageous IA | 0.108 | 0.097 | 1.230 | 0.267 | 1.11 | 0.92 – 1.35 |
| Advantageous IA | 0.089 | 0.093 | 0.894 | 0.344 | 1.09 | 0.91 – 1.31 |
| Prior expectation | 0.010 | 0.092 | 0.012 | 0.912 | 1.01 | 0.84 – 1.21 |

**Supplementary Table 4. Mixed-effects model with IQ and sex effects for the Coordination Game (N=202)**

Results of the binomial generalized linear mixed model (GLMM) testing confounding effects of IQ and sex on choice behaviour in the Coordination Game. Predictors of interest are underlined. GLMMs are described fully in the main text. Full R-code of this model is: mixed(Choice~IQ + sex + cAge-linear*Environment + cAge-quadratic*Environment+ cDisadvantageousIA*Environment + cAdvantageousIA*Environment + cpriorCoordinationGame*Environment + (Environment | Subject), method="LRT", family=binomial, data = dat, na.action=na.exclude, control=glmerControl(optimizer = "bobyqa", optCtrl = list(maxfun = 100000)))

| | Estimate | *SE* | $\chi^2$ | *P* | *Odds Ratio* | *CI* |
|---|---|---|---|---|---|---|
| Sex | -0.242 | 0.067 | 0.123 | 0.720 | 0.98 | 0.86 – 1.11 |
| IQ | -0.094 | 0.067 | 1.914 | 0.167 | 0.91 | 0.80 – 1.04 |
| Environment | -1.041 | 0.086 | 115.785 | <.001 | 0.35 | 0.30 – 0.42 |
| Age linear | -0.205 | 0.072 | 7.949 | 0.005 | 0.81 | 0.71 – 0.94 |
| Age quadratic | 0.121 | 0.068 | 3.114 | 0.078 | 1.13 | 0.99 – 1.29 |
| Disadvantageous IA | 0.160 | 0.070 | 5.125 | 0.024 | 1.17 | 1.02 – 1.35 |
| Advantageous IA | -0.168 | 0.066 | 6.281 | 0.012 | 0.85 | 0.74 – 0.96 |
| Prior expectations | 0.022 | 0.065 | 0.115 | 0.734 | 1.02 | 0.90 – 1.16 |
| Environment * Age linear | -0.445 | 0.095 | 21.122 | <.001 | 0.64 | 0.53 – 0.77 |
| Environment * Age quadratic | 0.062 | 0.090 | 0.466 | 0.495 | 1.06 | 0.89 – 1.27 |
| Environment * Disadvantageous IA | 0.092 | 0.090 | 1.022 | 0.312 | 1.10 | 0.92 – 1.31 |
| Environment * Advantageous IA | -0.256 | 0.086 | 8.503 | 0.004 | 0.77 | 0.65 – 0.92 |
| Environment * Prior expectation | 0.029 | 0.030 | 0.116 | 0.734 | 1.04 | 0.87 – 1.22 |

**Supplementary Table 5. Mixed-effects model for the Non-social learning task (N=245)**

Results of the binomial generalized linear mixed model (GLMM) testing effects of age (linear and quadratic) on choice behaviour in the Non-social Game. Significant effects are in bold.

| | Estimate | *SE* | $\chi^2$ | *P* | *Odds Ratio* | *CI* |
|---|---|---|---|---|---|---|
| **Age linear** | **0.251** | **0.082** | **9.150** | **0.002** | **1.29** | **1.09-1.51** |
| Age quadratic | 0.143 | 0.083 | 2.998 | 0.083 | 1.15 | 0.98-1.36 |

**Supplementary Table 6. Mixed-effects model with IQ and sex effects for the Non-social learning task (N=245)**

Results of the binomial generalized linear mixed model (GLMM) testing confounding effects of IQ and sex on choice behaviour in the Non-social learning task. Predictors of interest are underlined.

| | Estimate | *SE* | $\chi^2$ | *P* | *Odds Ratio* | *CI* |
|---|---|---|---|---|---|---|
| Age linear | 0.183 | 0.082 | 4.936 | 0.026 | 1.20 | 1.02-1.41 |
| Age quadratic | 0.114 | 0.080 | 2.027 | 0.155 | 1.12 | 0.96 -1.31 |
| IQ | 0.334 | 0.075 | 18.963 | <.001 | 1.40 | 1.20-1.62 |
| Sex | -0.074 | 0.078 | 0.892 | 0.345 | 0.93 | 0.80-1.08 |

**Supplementary Table 7. Parameters of best fitting computational models per game**

Table with all parameters of the best fitting computational models (i.e., models which include social preferences and decaying learning rates) per game and per age cohort. Results of the non-social game are also included.

**TRUST GAME**

| Age cohort | prior | learning rate $t$=0 ($\lambda_0$) | decay ($\tau$) | decision sensitivity ($\vartheta$) |
|---|---|---|---|---|
| 8-11 | 0.56 | 0.12 | 0.00 | 0.15 |
| 12-14 | 0.54 | 0.82 | 1.27 | 0.12 |
| 15-18 | 0.49 | 1.00 | 5.00 | 0.23 |
| 19-23 | 0.50 | 0.96 | 5.00 | 0.24 |

**COORDINATION GAME**

| Age cohort | prior | learning rate $t$=0 ($\lambda_0$) | decay ($\tau$) | decision sensitivity ($\vartheta$) |
|---|---|---|---|---|
| 8-11 | 0.71 | 0.67 | 1.94 | 0.63 |
| 12-14 | 0.67 | 0.61 | 1.31 | 0.46 |
| 15-18 | 0.65 | 1.00 | 5.00 | 0.53 |
| 19-23 | 0.68 | 1.00 | 5.00 | 0.69 |

**NON-SOCIAL LEARNING TASK**

| Age cohort | prior | learning rate $t$=0 ($\lambda_0$) | decay ($\tau$) | decision sensitivity ($\vartheta$) |
|---|---|---|---|---|
| 8-11 | 0.50 | 0.14 | 2.99 | 1.60 |
| 12-14 | 0.50 | 0.06 | 1.80 | 2.93 |
| 15-18 | 0.50 | 0.05 | 1.87 | 4.80 |
| 19-23 | 0.50 | 0.20 | 1.43 | 1.65 |

#### Participant instructions

Here we included the instructions for the Trust Game and the non-social learning task. Original instructions were in Dutch. The instructions for the Coordination Game are highly similar to instructions for the Trust Game, and are therefore not included here. Note that on almost every introduction screen for the behavioural tasks, figures of the task were included. Here the figures are only shown when necessary for understanding the accompanying text. The instructions also included some control questions; participants could only continue to the next screen when answered correctly. Full testing and instruction materials can be obtained from the corresponding author.

##### 1. Instructions Trust Game

--

The game you will play now has 3 parts. Each part has 30 short rounds. In each round you will make 1 choice.

On the next screens, the game will be explained.

--

In each round you play with another student from another school who also participated in this game.

In each round you will both make 1 choice.

In every round, the other is a **new** person.

--

The game will look like this:

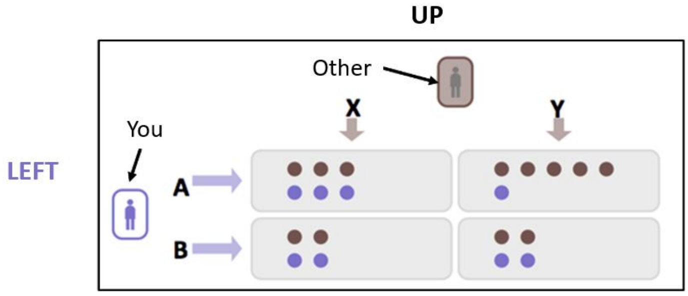


You are the **purple** icon on the left, and you can choose between the 2 purple arrows (**A** and **B**).

The other is shown on the top of the screen, and can choose between the 2 top arrows (**X** and **Y**).

--

In each of the four boxes you see dots. Each dot represents 1 point.

The points that you can win with your choice are always **purple**.

The points that the other player can win, have the other colour.

--

In every round you will make a choice between **A** and **B**.

The other will make a choice between **X** and **Y**.

Your choices **together** determine how many points you and the other win.

--

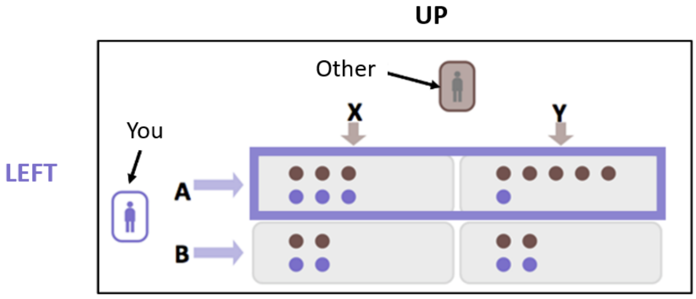


If you choose the **purple** arrow **A**, you select the two top boxes.

--

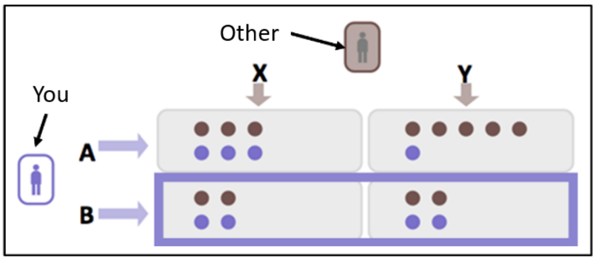


If you choose the **purple** arrow **B**, you select the two bottom boxes.

--

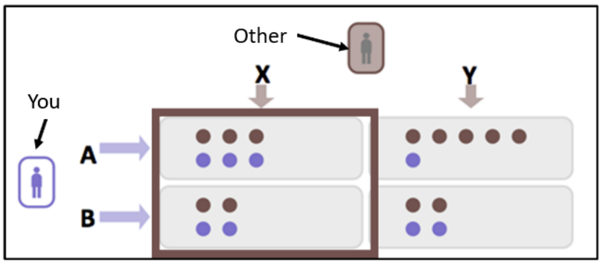


The other can choose arrow **X (top)** to select the two boxes on the left.

--

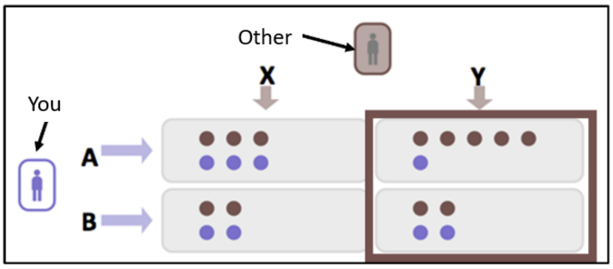


The other can choose arrow **Y (top)** to select the two boxes on the right.

--

The number of points **you** win, depends on your choice (**A** or **B**), but also on the *other player*'s choice (**X** or **Y**).

The number of points **the other player** wins, depends on the other player's choice (**X** or **Y**), but also on *your choice* (**A** or **B**).

Only *after* you submit your choice, you can see the other's choice.

--

For example:

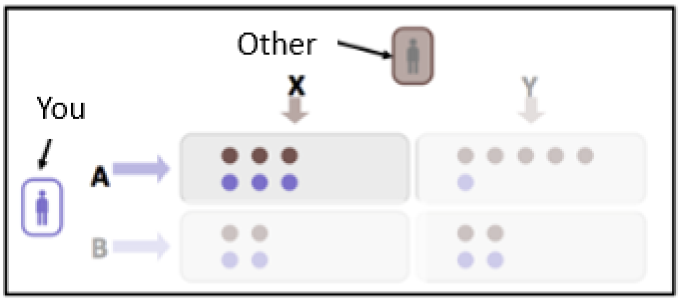


Suppose that you choose A, and the other chooses X:

- You would win **3 points**

- And the other would also win **3 points**

--

For example:

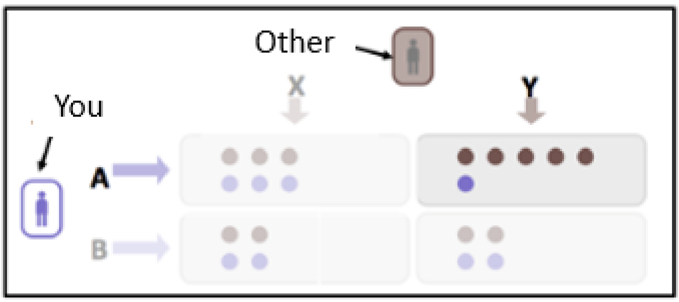


Suppose that you choose A, and the other chooses Y:

- You would win **1 point**

- And the other would win **5 points**

--

For example:

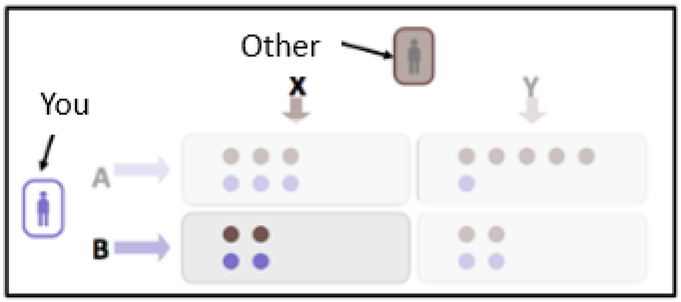


Suppose that you choose B, and the other chooses X:

- You would win **2 points**

- And the other would also win **2 points**

--

For example:

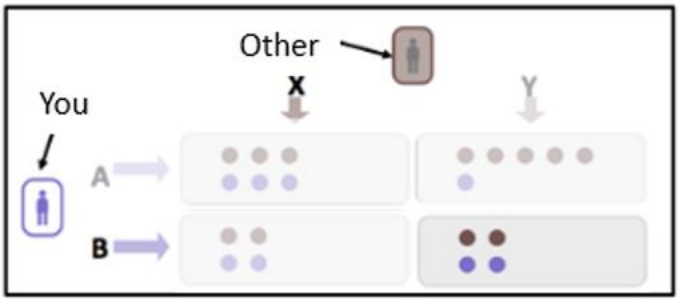


Suppose that you choose B, and the other chooses Y:

You would win ...... points (fill in)

And the other would win ...... points (fill in)

--

**That is correct!**

After you see what you and the other have won, the round is over and you will play with a **<u>new person</u>**.

**Important**: The choices of the other players are made previously by other students from another school.

--

At the end of the game, the computer will select 5 random rounds.
Each point in those 5 rounds is worth 1 lottery ticket.
After all games, we will draw 1 lottery ticket from all lottery tickets at your school.
The winner receives the gift voucher!
So the more points you have, the larger your chance to win.

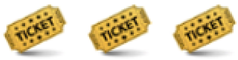


--

The other players can also earn lottery tickets with their choices. With the lottery tickets they also have a chance to win a gift voucher at their school.
Remember: your choices may affect the number of points **<u>you</u>** can win, but they may also affect the number of points the **<u>others</u>** can win.

--

**Environments**

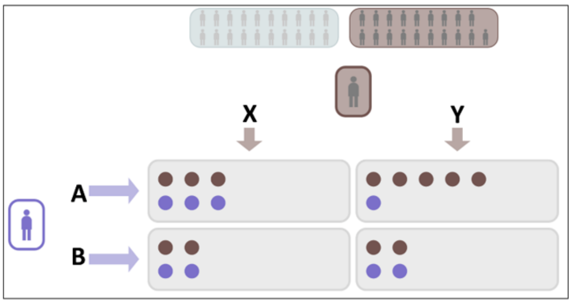


There are 2 environments of other players, each with their own colour.
The colour of the other player's icon tells you what environment they are in.

**In one environment, players usually choose <u>X</u> (left), and in the other environment players usually choose <u>Y</u> (right).**

--

In each round you will play with a new player from 1 of the 2 environments.

--

**Check Question**

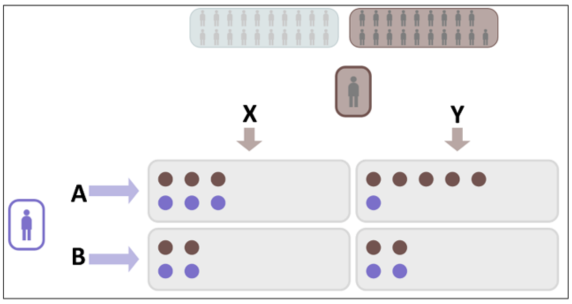


Suppose that you choose **B** and the other chooses **X**.
How many points would you win? (fill in)____

How many points would the other win? (fill in)____

--

**You have correctly answered the check question!**

There will now be 2 practice rounds before you start the real game.

Make your choice by clicking on 1 of your arrows. Then click 'Confirm' at the bottom of the page.

Next, you can see the other's choice, and how many points you and the other have won in this round.

Then click 'Continue' to start with the next round.

--

*{2 practice rounds}*

--

**Check question**

Which environment did the Other player belong to?

If you don't remember, please click back.

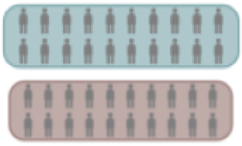

--

The game is about to start.

Do you have any questions? Please ask the researcher now.

You can click **start** if you understand the game.

**Try to win as many points as possible by paying close attention to what the other players are doing!**

--

**End of part 1**

You are finished with part 1 of the game!

--

#### 2. Instructions Non-social learning task

**Part 3**

In Parts 1 and 2 you were playing with other people.

In Part 3, you will not be playing with other people. Instead, you will play with computers.

--

The game will look like this:

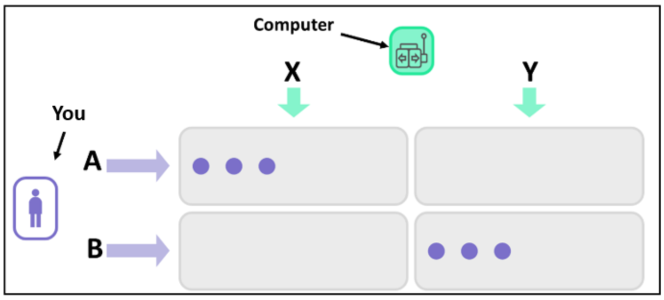


You are shown in purple on the left, and you can choose between the two purple arrows (A and B).

The computer can choose between the arrows on the top.

Your points are the purple dots.

--

As before, you will choose A or B (top or bottom boxes)

We have programmed the computer to choose X or Y (left or right boxes)

--

Your choice AND the computer's choice together determine how many points you will win.

**Important: the number of points you can win with your choice is different than in the previous games.**

--

For example,

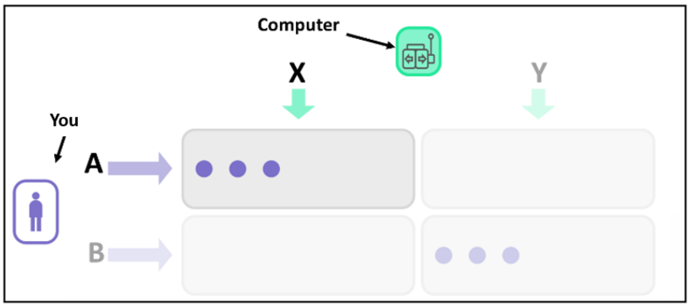


If you choose A and the computer chooses X:

You would win **3 points**

--

For example,

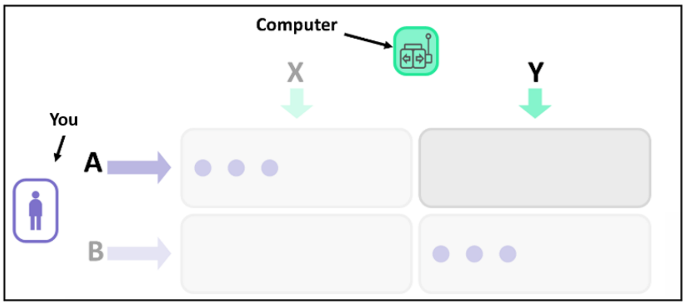


If you choose A and the computer chooses Y:

You would win **0 points**

--

For example,

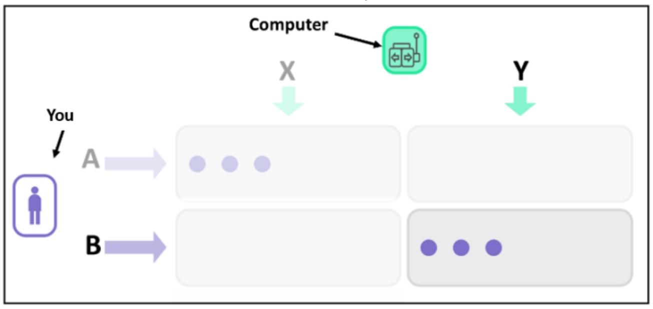


If you choose B and the computer chooses Y:

You would win **3 points**

--

For example,

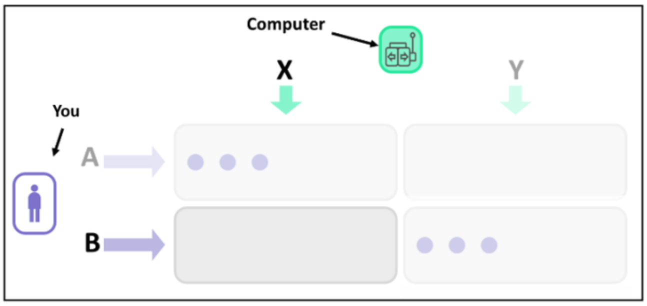


If you choose B and the computer chooses X:

You would win ____ **points** (fill in)

--

**That is correct!**

After you see what you have won, the round is over and you will play with a **new computer**.

--

At the end of the game, 5 rounds will be randomly selected.

Again, each point in the selected rounds is worth 1 lottery ticket.

Remember that you can win the gift voucher.

The more points you have the larger your chance of winning.

--

**Environments**

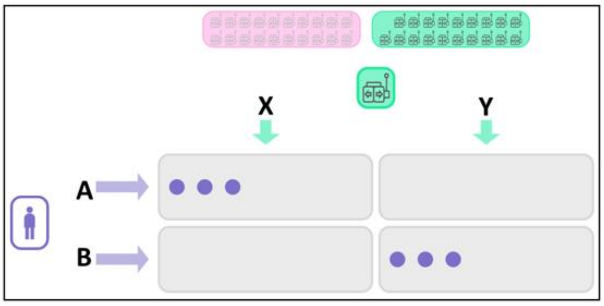


There are 2 environments of computers, each with their own colour.

The computer's colour tells you what environment the computer is in.

In one environment the computers usually choose **X** (left), and in the other environment the computers usually choose **Y** (right).

In each round you will play with a new computer from 1 of the 2 environments.

--

Click **Start** to start the game.

##### 3. Prior expectations

At the end of the learning task instructions, right before the start of the task, we asked about the prior expectations of the participants. This was only done for the social games (Trust Game and Coordination Game).

###### Prior expectations Trust Game

--

We are about to start the real game.

Before we start, we have a few questions.

--

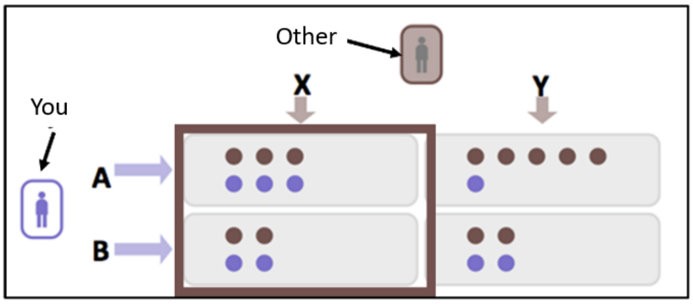


We are curious about what you think of the other players in this game.

Suppose that there are 10 other players, how many of these 10 do you think will choose X?

###### Prior expectations Coordination Game

--

We are about to start the real game.

Before we start, we have a few questions.

--

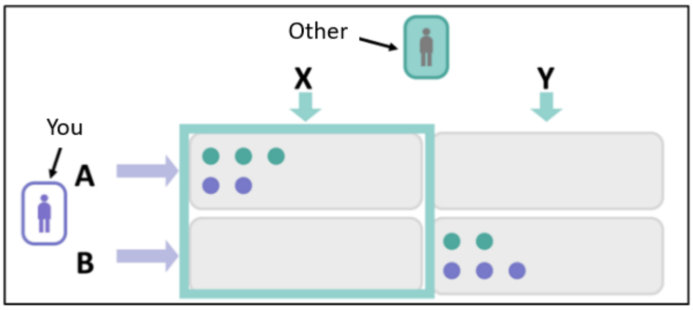


We are curious about what you think of the other players in this game.

Suppose that there are 10 other players, how many of these 10 do you think will choose X?
